# Interaction among bacterial cells triggers exit from lag phase

**DOI:** 10.1101/2022.01.24.477561

**Authors:** Maxime Ardré, Guilhem Doulcier, Naama Brenner, Paul B. Rainey

## Abstract

The relationship between the number of cells colonizing a new environment and time for resumption of growth is a subject of long-standing interest. In microbiology this is known as the “inoculum effect”. Its mechanistic basis is unclear with possible explanations ranging from the independent actions of individual cells, to collective actions of populations of cells. Progress requires precise measurement of lag time distributions while at the same time, experimentally controlling inoculum size. Here we use a millifluidic droplet device in which the growth dynamics of hundreds of populations founded by different numbers of *Pseudomonas fluorescens* cells, ranging from a single cell, to one thousand cells, were followed in real time. Our data show that lag phase decreases with inoculum size. The average decrease, variance across droplets, and distribution shapes, follow predictions of extreme value theory, where the inoculum lag time is determined by the minimum value sampled from the single-cell distribution. Our experimental results show that exit from lag phase depends on strong interactions among cells, consistent with a “leader-cell” triggering end of lag phase for the entire population.

## Introduction

When bacteria encounter new environmental conditions, growth typically follows four phases: a lag phase, during which bacteria acclimate, but do not divide; an exponential phase, during which cells multiply; a stationary phase, where nutrient exhaustion causes cessation of growth; and finally a death phase, during which cells may lyse.

In a fluctuating environment, each phase can play an important role in population persistence. The lag phase has particular significance because of both benefits (enhanced growth) and costs (sensitivity to external stressors) associated with the resumption of growth [16, 27]. Moreover, the time to resumption of growth — and controlling factors — has significant implications for the entire field of microbiology [15], but especially for infection caused by pathogens and for food safety [3, 19, 24].

Despite its discovery more than 100 years ago [17], cellular and molecular details defining the lag phase, factors triggering resumption of growth, and contributions to fitness, are not well understood. This is largely a consequence of the difficulties associated with experimental quantification of the dynamics of populations founded by small numbers of cells. Nonetheless, advances over the last decade have shown that bacteria in lag phase are transcriptionally and metabolically active [21], that lag phase is a dynamic state, that single cells are heterogeneous in time to resume division [11, 16, 27], and that numerous factors affect lag phase duration [3].

Arguably the most intriguing aspect of lag phase biology is the apparent inverse relationship between the number of cells in the founding population and duration of lag phase — often referred to as the “inoculum effect”. First reported in 1906 [20], the relationship has been shown to hold for a number of different bacteria [12–14, 18], although there exist few recent quantitative investigations. In certain instances, the inoculum effect is observed only under specific culture conditions [1, 12].

Factors controlling the inoculum effect are of special interest [1, 3, 6, 10, 13, 24]. Given that bacterial cells are typically variable in many of their properties, the simplest explanation posits that population lag time is determined by the set of cells with the shortest time to first division. Accordingly, the larger the founding population, the more likely it is that the inoculum contains cells on the verge of division, with these cells contributing disproportionately to the resumption of population growth.

An alternate explanation is that resumption of growth depends on interaction among founding cells, for example, via production of an endogenous growth factor: once a critical threshold concentration is achieved growth resumes, and the larger the inoculum the sooner this happens. Evidence in support of such control derives from analysis of *Bacillus* [13], *Francisella* (formerly *Pasturella*) *tularensis* [10], *Micrococcus luteus* [25] and *Aerobacter aerogenes* [6].

In instances where exit from lag-phase is determined by interactions among founding cells, models have assumed that all cells are equal contributors to the production of growth activating factors [12, 13]. However, an alternative possibility exists, namely, that population lag time is set by the activity of a single “leader cell” that triggers resumption of growth for the entire population of cells. Distinguishing among competing hypotheses requires precision measurements of population growth, high levels of replication, ability to control inoculum size, and crucially, knowledge of the distribution of lag times for single cells.

Here we use a millifluidic droplet device in which the growth dynamics of hundreds of populations founded by different numbers of *Pseudomonas fluorescens* cells were followed in real time. Our data confirm that lag phase shortens with inoculum size and provide a quantitative characterization of the effect: average decrease, as well as variance and distributions across repeated experiments for various controlled inoculum sizes. We demonstrate that these statistical properties follow extreme value theory, where population lag time is determined by the minimum value sampled from the single cell distribution. Additionally, we show that the inoculum effect cannot be explained by a selective sweep initiated from a small number of cells but rather involves the parallel growth of many lineages. These results suggest that exit from lag phase depends on strong interactions among cells, consistent with a leader-cell triggering end of lag phase for the entire population.

## Materials and Methods

### The strain

The ancestral strain of *Pseudomonas fluorescens* SBW25 was isolated from the leaf of a sugar beet plant at the University of Oxford farm (Wytham, Oxford, United Kingdom [22]). The strain was modified to incorporate, via chromosomally integrated Tn*7*, the gene GFP-mut3B controlled by an inducible Ptac promoter.

### Preparation of the cells

*P. fluorescens* SWB25 was grown in casamino acid medium (CAA). CAA for 1l: 5g of Bacto Casamino Acids Technical (BD ref 223120), 0.25g *MgSO*_4_ · 7 · *H*_2_*O* (Sigma CAS 10034-99-8), 0.9g *KH*_2_*PO*_4_ (Melford CAS 7758-11-4). Prior to generation of droplets SBW25 was grown from a glycerol stock for 19h in 5ml of CAA incubated at 28°C and shaken at 180 rpm. At 19h this culture was centrifuged at 3743 RCF for 4min and the supernatant removed from the pellet. The pellet was then resupended in 5ml of sterile CAA. It was then centrifugated and resuspended one more time. This prevents any interference of a growth-activator that may come from the supernatant of the overnight culture. The washed culture was then adjusted to OD 0.8 with CAA and mixed 1:1 in volume with autoclaved 30%v/v glycerol. 100μl aliquots were pipetted in 1ml eppendorf and frozen at −80°C. After freezing, one aliquot was taken to measure viable cells by plating on agar. We found 1.62 · 10^8^ cell·ml^−1^.

### Generation of droplets with a range of inoculum sizes

Each experiment with a range of inoculum sizes was prepared as follows. One frozen aliquot was thawed and diluted in 4ml of sterile CAA (with appropriate intermediate dilutions) by 7.04 · 10^4^×, 1.76 · 10^4^×, 1.76 · 10^4^×, 4.4 · 10^3^×, 1.1 · 10^3^×, 2.75 · 10^2^×, 6.875 · 10^1^×. We completed the dilutions from frozen stock by adding 29.1μl, 29μl, 28.6μl, 27.3μl, 21.8μl and 0μl, respectively, of sterile 60% v/v glycerol. This step is very import to balance the glycerol coming from the frozen stock and ensure that all the tubes have the same composition of medium (see Supplementary Fig. 5). We then added 50μl of sterile IPTG (100mM) to each sample. Each dilution was then pipetted in wells (250μl/well) of a 96 well microtitre plate to proceed to the generation of the droplets in the Millidrop Azur©. Droplets have a volume of 0.4μl which yields, with our dilutions, a range of inoculum sizes as follows: 1, 4, 16, 64, 256, 1024 cells/drop. We generated 40 replicate droplets for each population of a given founding inoculum size, except for populations founded by 1024 cells, which for technical reasons were restricted to 30 replicates.

### The inoculum of droplets follows a Poisson distribution

Importantly, the inoculum size in each droplet is controlled by the Poisson process occurring during the formation of a drop from the 96 well-plates. Therefore the inoculum that we report here is the average of the corresponding Poisson distribution. In particular the variance of the number of cells between the droplets is equal to the average inoculum.

### Generation of droplets with inoculum 1

To generate droplets with an inoculum of 1 cell/drop we diluted in sterile CAA a frozen alicot by 7.04 · 10^4^×, added 29.1μl of sterile 60% v/v glycerol and 50μl of sterile IPTG (100mM). We generated 230 droplets in the Millidrop Azur that yielded 156 droplets that grew due to the Poisson process inherent to the sampling process.

### Log-normality of the single cell lag time

The CDF for single cell lag times is displayed Fig. 2B, and is well described by a lognormal fit. To quantify the goodness of fit, we used the Shapiro–Wilk test with the null hypothesis that a sample log *θ*_1_, …, log *θ_n_* is derived from a normal distribution. The null hypothesis was tested with significance level alpha 5% and gave a p-value of 0.840 indicating that we can not reject the lognormality of the single cell lag time distribution. In addition, a quantile-to-quantile plot of single cell lag time against a lognormal distribution is shown in Fig. 11.

### Calculation of the uncertainty of measurement for the inoculum 1

Δ*N_th_* is the uncertainty on the threshold *N_th_* = 1.6 · 10^8^ cell·ml^−1^. The uncertainty of the calibration Fig. 12 gives Δ*N_th_* = 0.7 · 10^8^ cell·ml^−1^ for this value as depicted by the grey area. Δ*t_th_* is the uncertainty of the time when the population reaches beyond the threshold *N_th_*. We take its value as equal to the sampling frequency of the machine Δ*t_th_* = 18min. *λ* is the average growth rate of populations in droplets and Δ*λ* is the uncertainty. These quantities are estimated with the distribution of the growth rate shown Fig. 10: *λ* = 0.84 h^−1^ and Δ*λ* = 0.02 h^−1^. Δ*N*_0_ is the uncertainty on the inoculum size. In the experiment with inoculum 1 shown Fig. 2B, 230 − 156 = 74 droplets were empty despite being generated from the same mother culture. This is due to the randomness of the pipetting process that fills droplets of bacteria according to a Poisson process. The randomness of the process gives intrinsically an uncertainty on *N*_0_. In the following we explain how this was estimated. Knowing the number of empty droplets in the experiments allows calculation of the precise average inoculum of the experiment which correspond to the *λ* parameter of a Poisson distribution having its first value *p*(0) = *exp*(*−λ*) = 74/230 : *λ* = 1.134 cells/drop. This average takes into account the empty droplets with zero bacteria but we only measure the non-empty droplets. To estimate the average inoculum of the non-empty droplets we draw numerically a large series of random numbers with a Poisson probability of parameter *λ* = 1.134 and calculate the average and the SD of the non zero values. This yielded an average of 1.7 and an SD of 0.9. Therefore we consider that for our experiment in an ideal case with an infinite number of droplets the uncertainty on the inoculum of the droplets will be intrinsically Δ*N*_0_ = 0.9 cell/drop and that the averaged inoculum (of the filled droplets) is *N*_0_ = 1.7 cell/drop. All together these values allow calculation of the uncertainty of the lag time estimated by equation 1. The expression of uncertainty is given by equation 2 and numerical application gives Δ*θ* = 0.88 h.

## Results

### High-throughput quantification of bacterial population dynamics with millifluidic technology

To investigate the relationship between inoculum size and duration of lag phase, we used a millifluidic device to quantify the dynamics of bacterial population growth across time [2, 5]. The device allows the monitoring of 230 bacterial populations compartmentalized in droplets contained within a tube. Fig. 1A shows a portion of the tube with two droplets filled with cells. Statistical power of the experiment comes from precise control of large numbers of droplets, in terms of both inoculum size and homogeneity of culture conditions: Fig. 1B shows the growth dynamics of 40 replicate populations. Exponential growth and stationary phase are clearly seen, while lag time is concealed behind the detection threshold (grey area). Fig. 1C shows the growth dynamics of a population contained within a single droplet.

**Figure 1.**
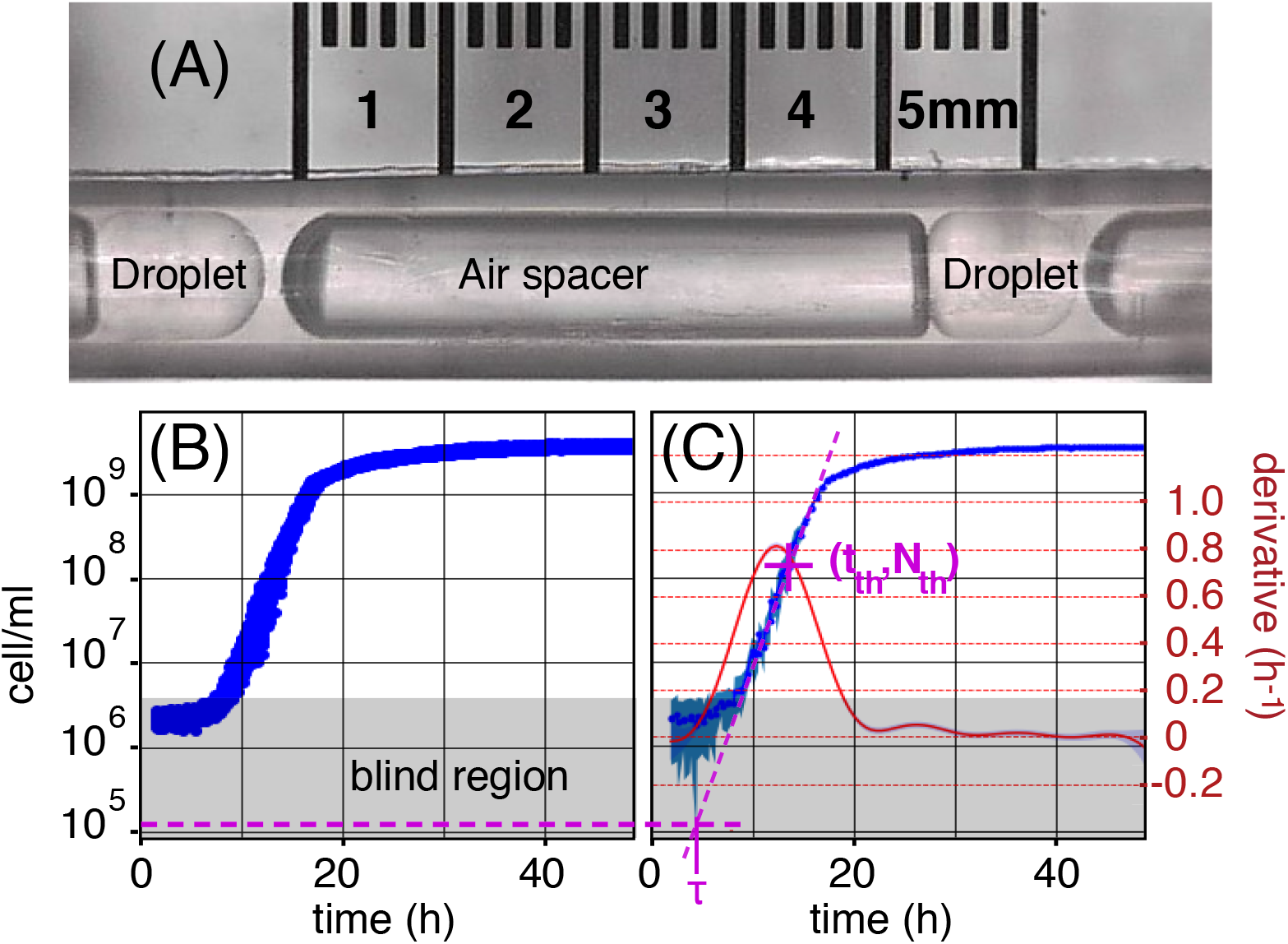
Bacterial population growth in droplets. **(A)** Two droplets of 0.4 μl are separated by an air spacer (to prevent droplet coalescence) inside the tube of a millifluidic machine. Droplets are prepared by “sipping” samples from a 96 well plate. Typically 230 droplets are produced from six mother cultures that differ solely in the number of founding cells (the *inoculum*). Each mother culture delivers 40 replicate droplets, but for technical reasons that last delivers 30 replicates. Droplets move back-and-forth, via changes in pressure, passing in front of a fluorescence detector every ~18 minutes. *P. fluorescens* SBW25 cells express GFP from a chromosomally integrated reporter, allowing changes in biomass to be determined based on intensity of the fluorescent signal (excitation at 497 nm emission at 527 nm). Signal intensity is calibrated to cell density by plate counting (Supplementary Fig. 12). The range of detection extends from 4 · 10^6^ to 5 · 10^9^ cell ml^−1^ (1.6 · 10^3^ to 2 · 10^6^ cells per droplet). The grey area in subfigures (B) & (C) denotes the region where bacterial density is below the threshold of detection. **(B)** Fluorescent signal across time from 40 replicate populations (in semi-logscale) in droplets prepared from the same mother culture. The average inoculum in each droplet is 1.6 · 10^5^ cell ml^−1^, or 64 cells per droplet. In this example the signal exceeds the detection threshold at ~7 h, by which populations are in exponential growth phase. At ~20 h, stationary phase is reached, marked by cessation of growth. **(C)** A single time series showing population growth within a single droplet coming from the set of replicates shown in (B). The left y-axis is shared between these two plots. The blue line depicts cell density derived using DropSignal [7] and the shaded area represents the standard deviation (SD). Population lag time is inferred as described in text. The purple dotted line crossing *N_th_* = 1.6 · 10^8^ cell ml^−1^ (64, 000 cells per droplet) extrapolates the exponential growth back to its intersection with the inoculum density (purple horizontal dotted line), giving *τ* ≈ 5 h. The red line gives the derivative of the time series, with shaded SD, and corresponding right y-axis in red.

**Figure 2.**
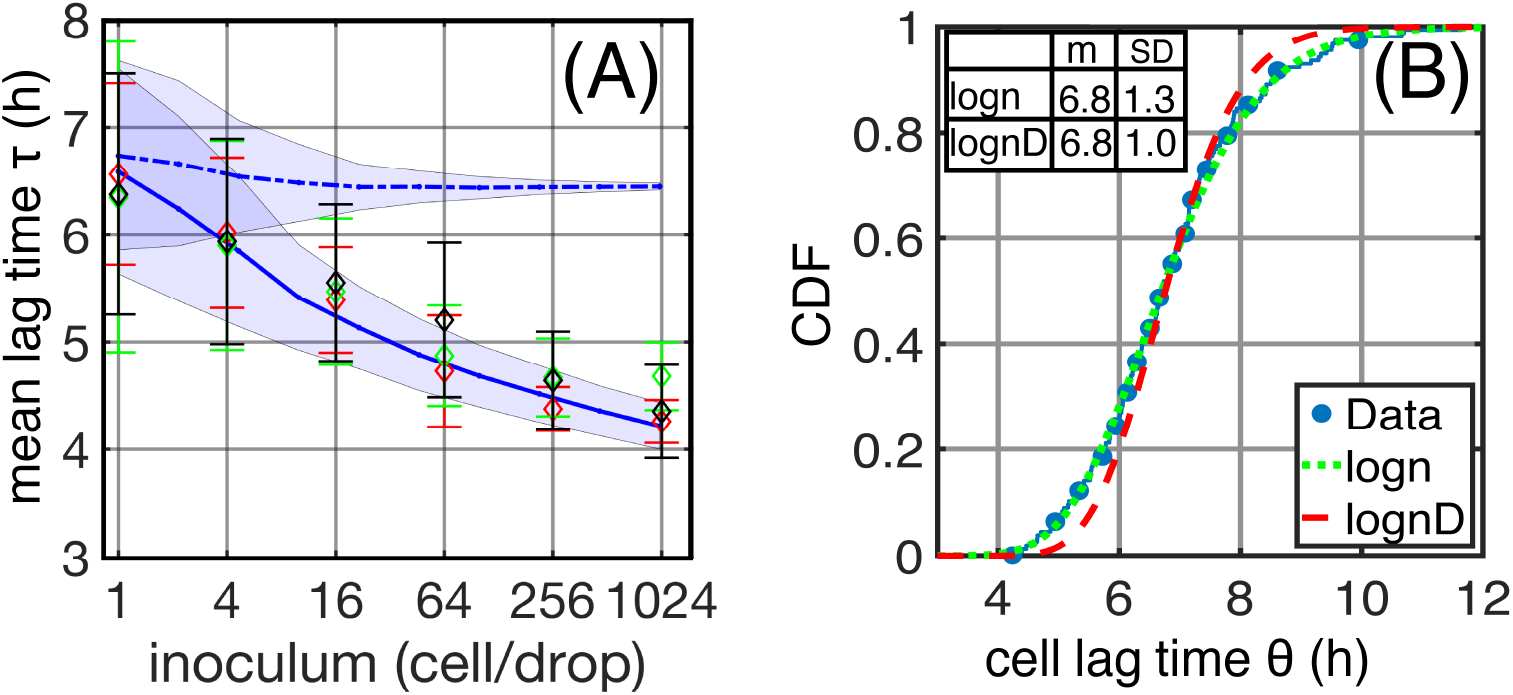
Quantitative data on lag-times are consistent with a strongly cooperative exit from lag phase. **(A)** Population lag time *τ* as a function of inoculum size for three independent experiments (colors). Diamond symbols are the mean lag times over several dozen droplets with a given inoculum size, with error bars denoting the standard deviation (SD). The data are compared to two models (blue lines - average, shaded blue - SD), based on single cell lag time heterogeneity: **(B)** Distribution of single cell lag times (*θ*) in an experiment where 156 droplets were inoculated with a single bacterium on average (blue points). CDF is the cumulative distribution function, where the y-value gives the probability that cell lag times assume a value less than or equal to the x-value. The measured distribution is fitted to a log-normal distribution (green dotted line) with a mean of 6.8h and a SD of 1.3h. A Gaussian “de-blurring” applied to these data generates the true distribution of cell lag times (red dotted line). Both models in (A) simulate populations founded by bacteria with lag times drawn at random from this corrected distribution. The dashed blue line shows the results for cells growing independently in the droplets. The continuous blue line shows results of a model where the lag time of all cells is set to the minimum in the inoculum, *i.e*., synchronization to leader cell.

Population density in the droplet is reported by fluorescence intensity from GFP-labelled bacteria with parameters describing the phases of growth being estimated from these time series (see Fig. 1C). Exponential growth rate, *λ*, is the maximum slope of the time series on a y-semi-logscale (we use a Gaussian processes method that makes no *a priori* assumption about the shape of the growth curve [23]). Final population size is estimated directly from measurements; death phase is not significant in our experiment and is ignored. Lag phase *τ*, is the time cells spend in a non-dividing phase prior to onset of exponential growth. Hardware limitations mean that fluorescence data are unobtainable for cell densities below 1,600 cells per droplet (4 · 10^6^ cell/ml) and thus *τ* must be estimated indirectly. This is done by firstly taking an arbitrary point (*t_th_, N_th_*) in exponential phase where cell density is *N_th_* = 1.6 · 10^8^ cell ml^−1^. By rearranging the equation for exponential growth: 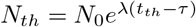, and making *τ* the subject

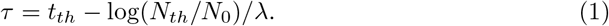

A geometrical counterpart of equation 1 gives the population lag time as the time point at which exponential growth (line in semi-log scale) intersects the horizontal line, which depicts inoculum density (see Fig. 1C). These measurements of lag-times provide a wealth of quantitative information on the inoculum effect, as described and interpreted in the following.

### Duration of lag phase depends on the number of founding cells

In Fig. 2A the average lag time from three independent experiments (colours) is shown as a function of inoculum size (diamonds). Lag time decreases monotonically from 6.4 ± 1.1h for droplets inoculated with a single cell, to 4.4 ± 0.3h for an inoculum size of 1,024 cells. The standard deviation (SD), represented by the error-bars in the figure, decreases monotonically and slowly with increasing inoculum size.

We also examined the dependence of other growth parameters on inoculum size. Initial experiments showed an effect on final cell density, however this was found to be a consequence of subtle differences in glycerol concentrations arising from dilutions of frozen glycerol-saline stock cultures used to prepare founding inocula. When corrected, no effect of inoculum size on final cell density was observed. This technical, but important experimental observation is explained in Supplementary Fig. 5. Additionally, no effect of inoculum size on mean growth rate was detected, although the variance across droplets decreased. Details are provided in Supplementary Fig. 8.

What might be the basis of the decrease in mean lag time with inoculum size? There are three mutually exclusive explanations, all recognise that populations of cells are heterogeneous with regard to individual cell lag times as a consequence of innate phenotypic variability. Explanation I posits no interaction among cells, with population lag time being set by an event equivalent to a selective sweep that is initiated by the cell (or cells) with the shortest cell lag time.

Explanations II and III involve interactions among cells and can be thought of in terms of two extremes of a continuum. Explanation II posits that all cells contribute equally to the production of some growth-stimulatory factor. Explanation III recognises that population lag time could be set by the cell with the shortest lag time and whose activity triggers division of other cells. We demonstrate below that distinguishing between these alternate explanations is possible via quantitative data obtained from the millifluidic droplet device. Making this distinction requires knowledge of the lag time distribution of populations founded by single cells.

### Precise estimation of the distribution of cell lag times from inocula containing a single bacterium

To quantify the lag time of individual bacteria, 230 droplets were inoculated by – on average – a single bacterium, resulting in growth in 156 droplets (the inoculation of droplets follows a Poisson process). For each droplet, the lag time was estimated as in Fig. 1C. The resulting distribution of lag times is shown in Fig. 2B (blue dots). In this case, the lag time of each population is clearly equal to that of the founding cell. We denote the single cell lag by *θ* to distinguish it from *τ* of larger inoculum size that may be affected by cooperative effects. The heterogeneity of cell lag times is broad, ranging from ~4 h to ~12 h, with a mean value of *m_exp_* = 6.8 h with SD *σ_exp_* = 1.3 h. A

Shapiro-Wilk test applied on the logarithm of the data reveals the underlying distribution to be log-normal (see also the quantile-to-quantile plot shown in Supplementary Fig 11). Fitting a log-normal function (green dashed line in Fig. 2) yields parameters *μ* = 1.9 and *s* = 0.2.

Although the fit is good, there is uncertainty in the estimation of lag times due to measurement errors that propagate to the extrapolation Eq. 1. This equation expresses the dependence of lag time on parameters *t_th_*, *N*_0_, *N_th_* and *λ* for droplet populations, including the special case of a population being founded by a single cell. Expanding it to a Taylor series and assuming independent variables allows the uncertainty Δ*θ* to be calculated as:

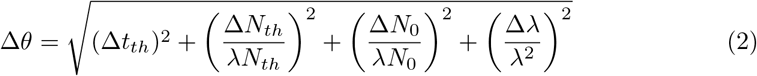

where Δ*t_th_*, Δ*N_th_*, Δ*N*_0_ and Δ*λ* correspond to the uncertainty of *t_th_*, *N_th_*, *N*_0_ and *λ*, respectively. Given the values of these uncertainties, we estimate Δ*θ* = 0.88h (see Materials and Methods for details of calculations).

The uncertainty associated with direct measurements blurs the “true” distribution of single cell lag times, which is less dispersed, *i.e*., has a smaller SD. We assume a Gaussian noise of zero mean and a SD equal to the measurement uncertainty *σ_noise_* = Δ*θ*. Deconvolution of the Gaussian noise from the measured distribution [4] amounts to subtracting its mean and variance from that of the measurements:

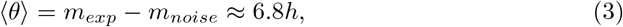

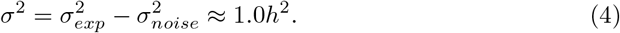

The corrected distribution remains lognormal, with parameters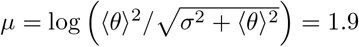 and *s*^2^ = log *σ*^2^/⟨*θ*⟩^2^ + 1 = 0.14. Expression of the true probability density of lag time is thus:

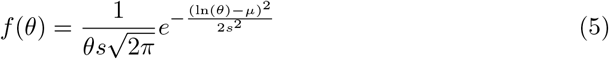

The red dotted line in Fig. 2B depicts the corresponding CDF after correction for measurement noise; it is narrower than that obtained by direct measurement. This distribution can now be used to examine the previously proposed explanations for the dependence of population lag time on inoculum size.

### A selective sweep initiated by cells with the shortest lag time is inconsistent with the data

Intuitively, one might imagine that the large variability in single-cell lag-times is sufficient to account for the observed inoculum effect, even for independently growing cells: larger inocula contain out-lier cells that are fast to resume growth; these could in principle take over the population and reach maximal cell number faster, causing the observed inoculum effect. Having a precise estimate of the single cell lag time distribution, it is now possible to put this hypothesis (Explanation I) to quantitative testing.

To this end, growth of populations within droplets established from different numbers of founding cells were simulated and the match with experimental data determined. Virtual droplets were founded by cells with lag times drawn from the true distribution (shown in Fig. 2B) and with exponential growth rates drawn from the measured distribution (see Supplementary Fig. 10). Cells were then allowed to replicate within droplets. To mimic the experimental protocol, the time *t_th_* at which populations reach *N_th_* = 64, 000 cells (equal to a density of 1.6 · 10^8^ cell·ml^−1^) was determined. Eq. 1 was then used to calculate the lag time of each population with known *N*_0_ and with known mean growth rate *λ*. The blue dotted line in Fig. 2A shows the results of these simulations.

In marked contrast to the experimental results, these simulations of independent (non-interacting) cells show almost no dependence of the mean population lag time on inoculum size. In addition, the decrease in variation across droplets, represented by the SD of lag time (shaded blue region around dotted line), decreases rapidly, whereas in the experimental data the SD decreases much slower.

Failure of explanation I to account for the data can also be understood by a simple calculation based on the growth characteristics. The corrected CDF of the single cell-lag times, Fig. 2B, has a value of 0.025 for lag time 5 h. In other words in a droplet inoculated by 1024 cells, ≈ 25 cells (0.025 × 1024) have a lag time equal to or shorter than 5h. Similarly, the number of cells exiting lag phase between 5.8 h and 7.8 h (around the mean 6.8 ± 1 h) is: (0.86 − 0.14) × 1024 ≈ 737 cells. Given the single cell growth rate *λ* = 0.84 h^−1^ (SI Fig. 10), the generation time is *g* = 0.83 h. Thus, the 25 cells that start dividing before 5 h go through roughly (6.8 − 5.0)/*g* ~ 2 generations before the 737 cells around the mean start dividing. Two generations of 25 cells yields 100 offspring; therefore, clearly cells with a short cell lag time do not exert a dominant sweep effect on the population.

The above estimate implies that, during the time of the measured growth, many lineages within droplets grow simultaneously and produce offspring. Therefore, if cells are independent, the population lag time is expected to be roughly equal to the mean of the single-cell lag time distribution, which is independent of inoculum size as the simulation shows. Moreover, as an av√erage over many cells, the lag time variability across droplets should decrease as 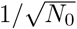 with inoculum size. Indeed, the SD of the simulation shown in Fig 2A decreases rapidly with inoculum size, in marked contrast to the experimental data – which decrease slowly. This discrepancy of the variance behavior with inoculum size indicates that population lag time does not arise as an average over independent cells in the inoculum. Alternative scenarios where cells are not independent are considered below.

### A leader cell triggering end of lag time for the population is consistent with the data

We now turn to test Explanations II and III that involve interactions among cells within the founding inoculum. At one extreme case (Explanation III), collective growth is governed by a single event that synchronises population growth to the cell with the shortest lag time. This would happen if the cell that divides first signals this effect to other cells, such that the entire population exits lag phase almost simultaneously. We first examine the consequences of this assumption and compare it to the data, and then consider the alternative scenario, namely, Explanation II, in which interactions among cells involve all cells contributing equally to the production of a growth-stimulating factor.

In statistical terms, we assume that an inoculum of *N*_0_ cells is a random sample from the single cell lag time distribution *f* (*θ*). If there is a leader cell that triggers growth for all other cells, the measured population lag time will be equal to the shortest cell lag time in the sample, *θ_min_*. Extreme Value Theory (EVT) provides a framework for statistical analysis of the extreme value of a sample, such as the shortest lag time *θ_min_* among *N*_0_ cells [8]. In the limit of large samples, EVT predicts the dependence of the mean and variance of a collection of *θ_min_* coming from samples of size *N*_0_. It also predicts that the distribution of *θ_min_* from populations founded by cells of a given inoculum size will approach a limiting fixed shape after appropriate normalization as the sample size increases; the precise shape is determined by global properties of the single cell distribution *f* (*θ*).

The unique features of our experiment create an ensemble of droplets with controlled inoculum size, and a measurement of the population lag time for each, labelled *τ*. These data provide the statistical properties required to test the hypothesis that *τ* = *θ_min_*(*N*_0_), namely that the population lag time is equal to the minimum cell lag time among the *N*_0_ single cells of the inoculum. For this we use predictions given by EVT on the distribution of *θ_min_* and ask whether they are consistent with the statistical properties of *τ* as measured in the droplets.

The first prediction is that both the average and the SD of *θ_min_* from populations decrease slowly with inoculum size *N*_0_. The precise scaling is derived from the single cell distribution *f* (*θ*) (see Appendix in the Supplementary Information); for a lognormal distribution we find the scaling to be:

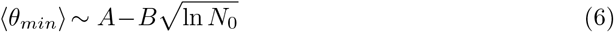

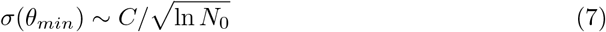

where *A, B, C* are constants. Both predictions agree well with the population lag time *τ*. Fitting the curve of Eq. 6 to the experimental data on the relationship between mean population lag time (*τ*) and inoculum size, reveals a close match (Fig. 3A&B). The same holds for the SD fitted to Eq. 7. We note that although testing this prediction involves fitting constants, the dependence on sample size *N*_0_ through 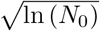 is nontrivial and unique to the predictions of EVT.

**Figure 3.**
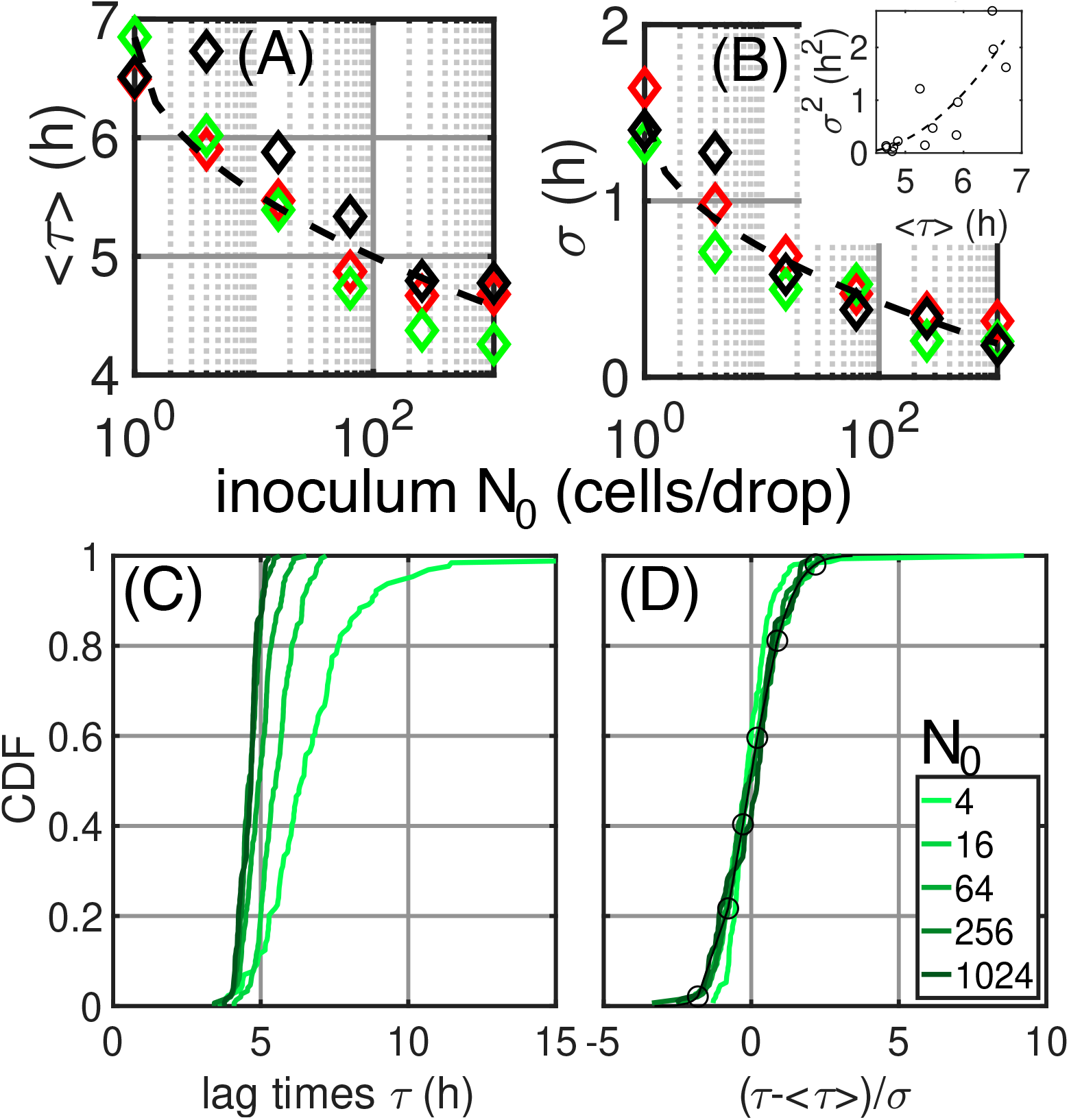
Statistical properties of lag times. For three independent experiments (coloured symbols), mean lag times **(A)** over populations and their SD **(B)** are depicted as a function of inoculum size. Inset: variance as a function of mean. The scaling relations predicted by EVT are shown in dashed lines: 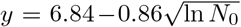 for the mean and 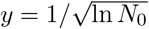 for the SD. Each point is calculated over 40 replicate populations (droplets). See Supplementary Appendix for details. **(C)** Cumulative Distributions of population lag times for different inoculum sizes (*N*_0_, colors; legend in (D)). The curves derive from the pooled data of three independent experiments yielding at least 120 population lag times for each. **(D)** Same data as in (C), scaled by subtraction of empirical mean and division by SD. The black line with circle markers depicts the fit by the universal distribution predicted by the EVT. The y-axis is shared between (C) and (D).

A further prediction is that the distribution of minimal values (*θ_min_*), drawn from different sample sizes, tends to a universal shape in the limit of large samples. Although strictly speaking this holds in the limit *N*_0_ → ∞, in practice it may be expected to hold also for finite samples - even relatively small ones. For each sample size *N*_0_, our experiment provides a distribution of population lag times (*τ*), estimated over all droplets with the same inoculum size. These CDFs are depicted in Fig. 3C for all inoculum sizes that are above 4 cells. To test whether the prediction on *θ_min_* holds for the population lag time *τ*, we normalize each CDF of *τ* by subtracting its mean and dividing its SD. Fig. 3D shows the result of this normalization and demonstrates that the distributions of *τ* collapse on one another, consistent with our hypothesis *τ* = *θ_min_*. The lightest shaded curve, corresponding to inoculum *N*_0_ = 4, deviates from the rest - possibly indicating that this sample size is too small to acquire the limiting shape.

The universal distribution itself is also predicted by EVT. Its CDF has the form:

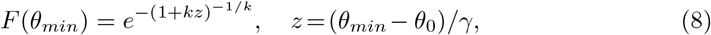

with location and scale parameters *θ*_0_, *γ* and a shape parameter *k* that reflects properties of the parent distribution *f* (*θ*), specifically the decay at its tails. Fitting the normalized data with this formula reveals an excellent match between the universal distribution formula (black line with circle marker in Fig. 3B) and the normalized measured distributions of *τ* (green lines), at least for inoculum sizes above 4 cells per droplet. The analytical formula for the distribution justifies the empirical procedure of normalizing by sample mean and standard deviation used above as a test for the universal shape (see Appendix in Supplementary Information).

As a corollary of the predictions in Eqs. 6,7, the variance and mean of the distributions of extreme values drawn from different sample sizes follow a well-defined relationship (see Appendix). The agreement of this relationship with the population lag time data is shown in the inset of Fig. 3B.

Taken together, the scaling of the mean and SD of *τ* according to the inoculum size (Eq. 6 & 7), the resulting relationship between variance and mean of *τ*, the collapse of normalized distributions of *τ* at different sample (inoculum) sizes, and the fit of the normalized distribution to the theoretical formula Eq. 8, are consistent with population lag time being equal to the shortest cell lag time in the inoculum: *τ* = *θ_min_*. With our understanding that growth involves multiple simultaneous lineages, we conclude that, at the time of the shortest lag time in the inoculum, many cells must start growing in parallel.

### A single leader cell determines population lag time

The agreement of statistical properties with predictions from EVT suggest that exit from lag phase is triggered by a single event – possibly a single leader cell – that signals the exit from lag to all other cells. To test this hypothesis we performed a set of simulations where cells are not independent. As for previous simulations, at each inoculum size thousands of virtual populations were generated (see simulation code in the Appendix of Supplementary Information). For each population the cell lag time of each founder cell was drawn at random from the experimental single cell lag time distribution (Fig. 2B) and then set to the shortest cell lag time in the sample. This means that all founder cells begin to proliferate at the same time as the leader cell with population lag time *τ* = *θ_min_*. As in the previous simulation, the numerical population was allowed to grow exponentially with population lag time being estimated as per the experiments.

The results are depicted by the solid blue line in Fig. 2A and are a close match to the experimental data over three orders of magnitude in inoculum size. Note that this is not a fit: the only input is the true single cell lag distribution measured for populations founded by one cells (Fig. 2B). Additionally, results from the simulations match the slow decrease of lag time variability among droplets observed in the experiments (shaded area Fig. 2A). Our results thus provide an explanation for the relationship between size of the founding inoculum and population lag time and are fully consistent with resumption of growth of all the cells of the inoculum triggered by an event effected by a leader cell.

Thus far it is not possible to be certain as to whether population growth is triggered by a single cell, or a small pool of cells, each having a short lag time. To further investigate we performed a further simulation where the fraction of cells contributing to start of growth varies continuously. In this simulation we consider that cells produce a growth activator which triggers growth of all cells in the inoculum upon passing a threshold. The concentration threshold effectively determines the fraction of cells in the inoculum that affect exit from lag phase. If the threshold is low, it is sufficient that a single cell produces the growth activator. If the threshold is set high, then multiple cells can end lag phase before the threshold is reached and contribute to the production of the activator. The duration of lag phase for every cell, is as before, set by drawing a random value from the experimental cell lag distribution Fig. 2B (see code in Supplementary appendix).

Running simulations for a range of activator thresholds at different inoculum sizes allows investigation of the number of cells that have ended lag phase before the occurrence of the triggering event. The results are depicted in the Supplementary Information Figs 6 and 7. First, it is seen in Fig. 6 that a strong dependence of population lag time on inoculum size appears only when the threshold concentration of the growth activator is low. Given the significant inoculum effect observed in our experiments, we conclude that this model is consistent with the data only in the region of low threshold. Second, Fig. 7 shows the effective number of cells that “lead” the population to exit from lag phase. This number is defined by those cells which have already reached their lag time as drawn from the single-cell distribution before the threshold was reached. Strikingly, we see that for a large range of activator concentrations, only a single cell needs exit lag phase in order to trigger growth of all cells in the population. This leads us to conclude that our experimental observations are fully consistent with Explanation III, where a single leader cell end lag phase for the entire population.

## Discussion

An inverse relationship between the number of bacterial cells founding a new environment and the time to exit lag phase was first noted more than 100 years ago [18, 20]. Despite its significance, rigorous validation has been lacking, and understanding of the causes and controlling factors remain incomplete. Lack of progress has stemmed largely from difficulties associated with experimental analysis of populations founded by few cells. Progress requires ability to follow the growth dynamics of replicate populations established from precisely determined numbers of founding cells. Additionally required is knowledge of the distribution of lag times for hundreds of single cells. Once known, statistical approaches can be used to link the distribution of population lag times, to the behaviour of single cells. The nature of the relationship stands to shed insight on cell-level causes of the inoculum effect.

Here, taking advantage of new opportunities presented by millifluidic technologies [2, 5] we have obtained quantitative evidence from highly replicated populations founded by controlled numbers of cells, that in populations of *P. fluorescens* SBW25, the time to resume growth after transfer to a new environment is strongly influenced by size of the founding inoculum. Moreover, the same technology has allowed determination of the duration of lag phase for a sample of individual cells. The average decrease in time to growth resumption, variance across droplets, and distribution shapes, follow predictions of extreme value theory, consistent with the inoculum lag-time being determined by the minimum value sampled from the single-cell distribution. At the same time, within droplets, growth of multiple cell lineages in parallel contribute to population expansion with no single lineage providing a disproportionate effect on duration of lag time. Our experimental results thus show that exit from lag phase depends on strong interactions among cells, consistent with a “leader cell” triggering end of lag phase for the entire population.

This finding builds on recent work in which the time to first division of single bacteria maintained in isolated cavities of microfluidic devices has been measured [11, 16, 27]. From such studies it is clear that there is substantial variation in cell-level lag time with evidence that this variance can have profound fitness consequences for population growth. For example, in fluctuating environments, heterogeneity in the time for individual cells to resume growth, can facilitate survival in the face of environmental change [11], especially that wrought by periodic antibiotic stress [9, 16, 27].

Although microfluidic chambers used for analysis of isolated cells allow precision measures of the distribution of lag times for single cells, such experimental devices do not allow for interactions among cells, thus making problematic any attempt to connect the distribution of single cell lag times to population lag times. In fact, extrapolation of population lag times from knowledge of the distribution of single cell lag times would be justified only in the case of independent cells.

Linking cell and population level behaviours necessarily requires measures of lag times both for individual cells and for populations in precisely the same environment. Moreover, the environment should be well mixed (spatially homogeneous and devoid of surface effects) so that should emergent population-level behaviours be relevant, mediated via, for example, production of diffusible growth factors [12, 13], then effects can be observed. In this regard the millifluidic device has proven fit for purpose.

In seeking an explanation for the observed inoculum effect we considered three mutually exclusive explanations. Central to Explanation I was absence of interactions among cells with the inoculum effect being explained by disproportionate growth of the set of cells with the shortest time to first cell division. Both simulations and simple calculations led to unequivocal rejection of this hypothesis.

Explanations II and III recognised the possibility of interactions among cells. Because of the power of extreme value theory combined with well understood statistical properties we chose to focus on whether population lag times were determined by the minimum value sampled from the single-cell distribution (Explanation III). EVT makes predictions as to the distribution of minimal cell lag times across droplets, which surprisingly, also held for the distribution of population lag times measured in the experiments, leading to the conclusion that the data are consistent with the population lag time being equal to the minimal cell lag time present in the inoculum. Simulations of population growth in droplets based on this evidence delivered an almost perfect match to experimental data. While conformity to Explanation III means that Explanation II in a strict sense (in which all cells contribute equally to exit from lag phase) cannot hold, the fact that there is a continuum of possibilities led us to perform additional simulations to address whether our data are consistent with resumption of growth being triggered by just a single leader cell. The findings fit surprisingly well and are fully consistent with a single cell being sufficient to trigger resumption of growth for the entire population.

An obvious next question concerns identity of the growth activator. While detailed investigations are beyond the scope of this study, we nonetheless, considered the possibility that iron chelation might trigger the effect. Such a possibility has been previously suggested [12]. To this end we repeated the initial experiment in which the time to resumption of growth was determined in replicate populations founded by different numbers of cells as in Fig 2A. Instead of SBW25 a mutant deficient in production of the iron chelating compound pyoverdin was used *P. fluorescens* SBW25 *pvdS* G229A [26]. No change in the inoculum effect was observed (see Supplementary Fig 15), thus ruling out pyoverdin as the activating molecule.

The relationship between the number of cells founding growth in a new environment and duration of lag phase has profound implications for microbiology. While much remains to be understood, including generality and molecular bases, the rigorous quantification achieved here via a millifluidic device provides unequivocal proof of an inoculum effect in *P. fluorescens* SBW25 and its cooperative nature. Moreover our statistical analysis of the distribution of population lag times is consistent with the activity of a single leader cell triggering simultaneous growth for all cells in the founding population.

## Acknowledgments

We thank Millidrop for development of the prototype Azur3 and their willingness to engage in active collaboration. Discussion and input from Arjan De Visser, Jerome Bibette, Andrew Farr, Lukas Geyrhofer, Isabelle Rivals, Clara Moreno-Fenoll, Jean Baudry, Nicolas Bremond, Jean-Baptiste Dupin, Wilfried Sire, Ankur Chaurasia is gratefully acknowledged. The work was supported by an HFSP grant “Interrogating bacterial social interactions in droplets” RGP0010/2015

## Author contributions

M.A. & P.B.R. designed the experiment and determined the strains and relevant experimental conditions, M.A. performed the experiments, M. A. & G.D. developed the software to analyse the data, M.A & N. B. made the modeling and simulation, M.A., N.B., P.B.R. wrote the paper.

## Supplementary Appendix

### Codes of simulation

First we give the code of the simulation running in Matlab©to generate the Figs. 2A and 6&7:

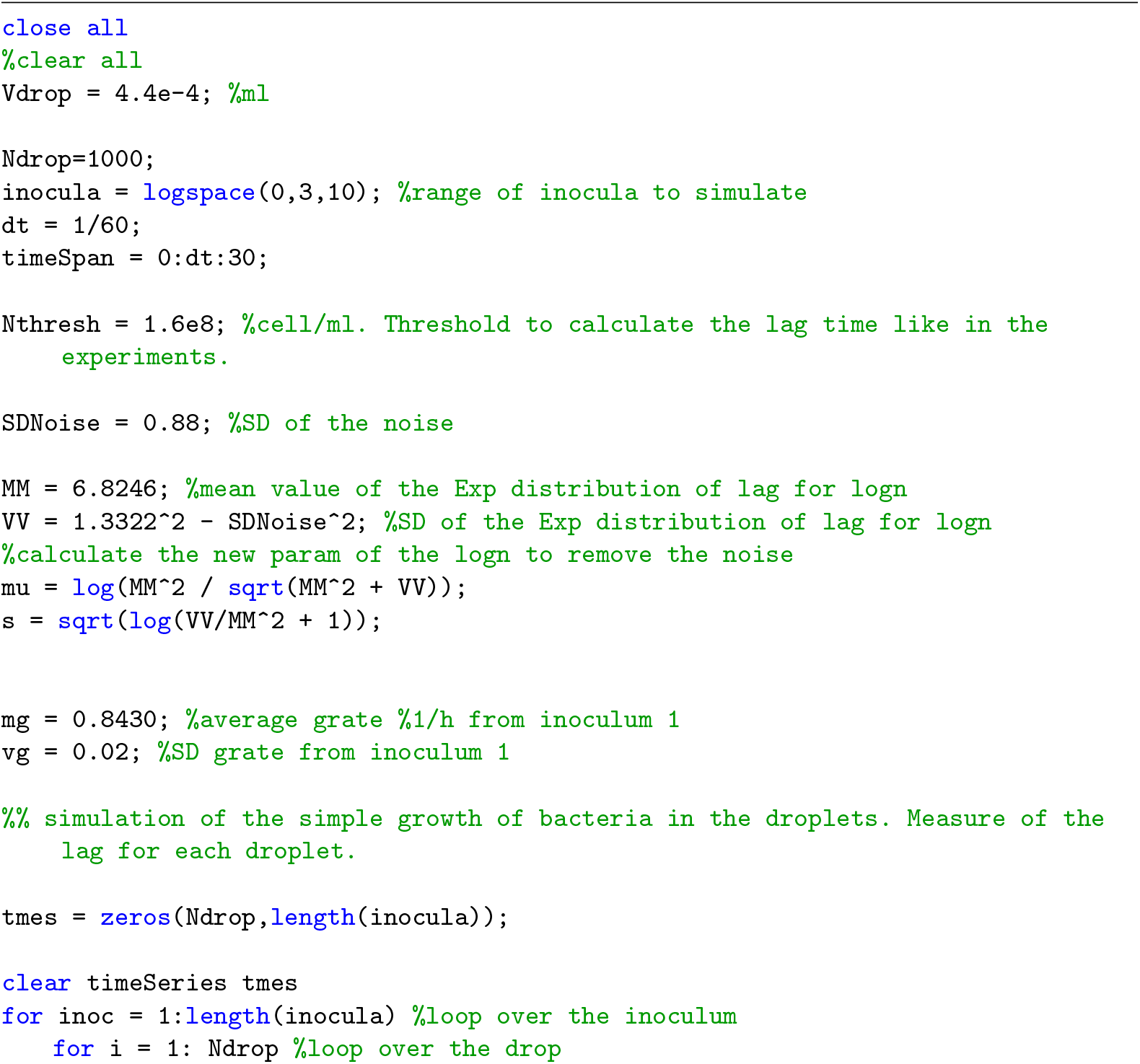
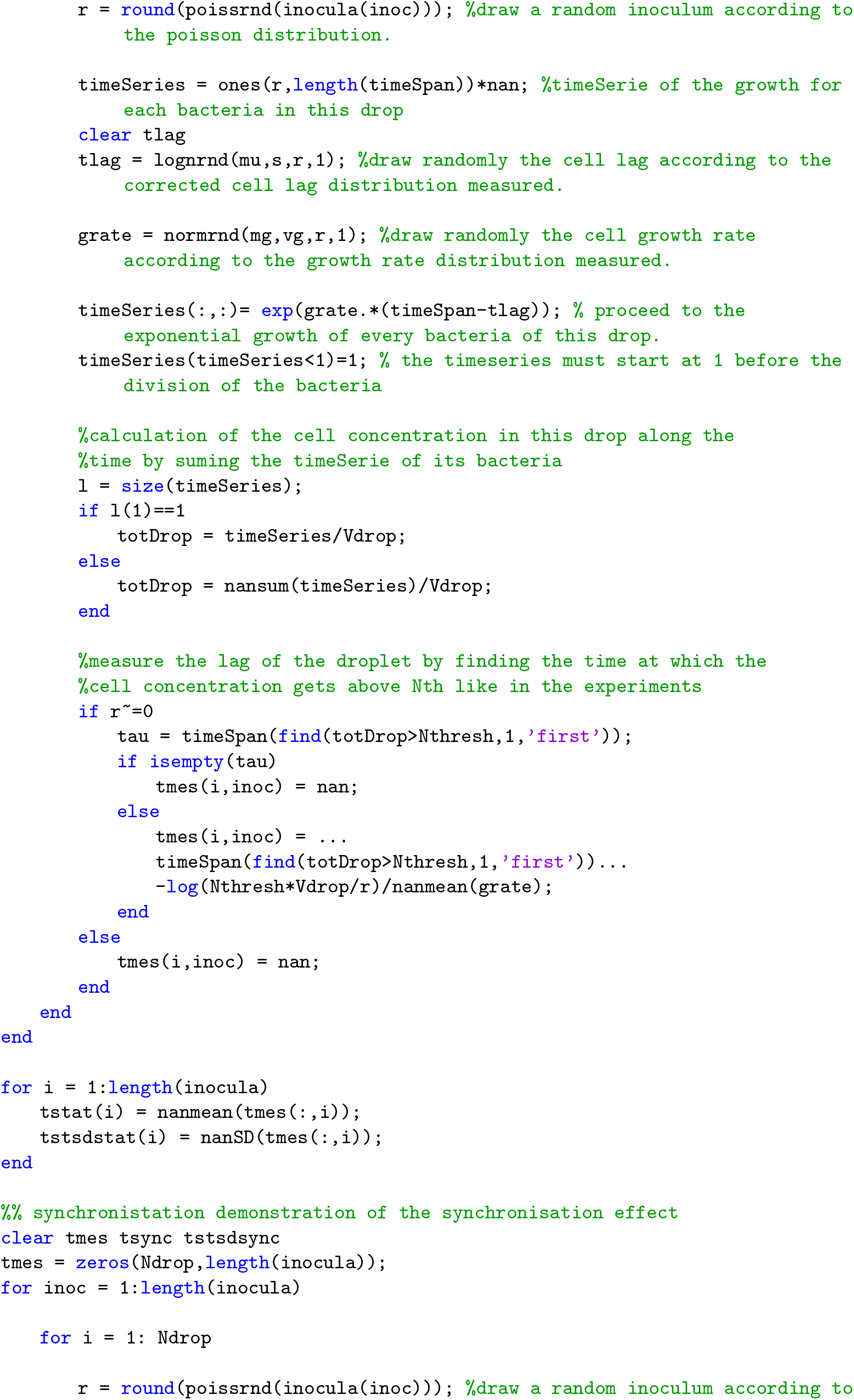
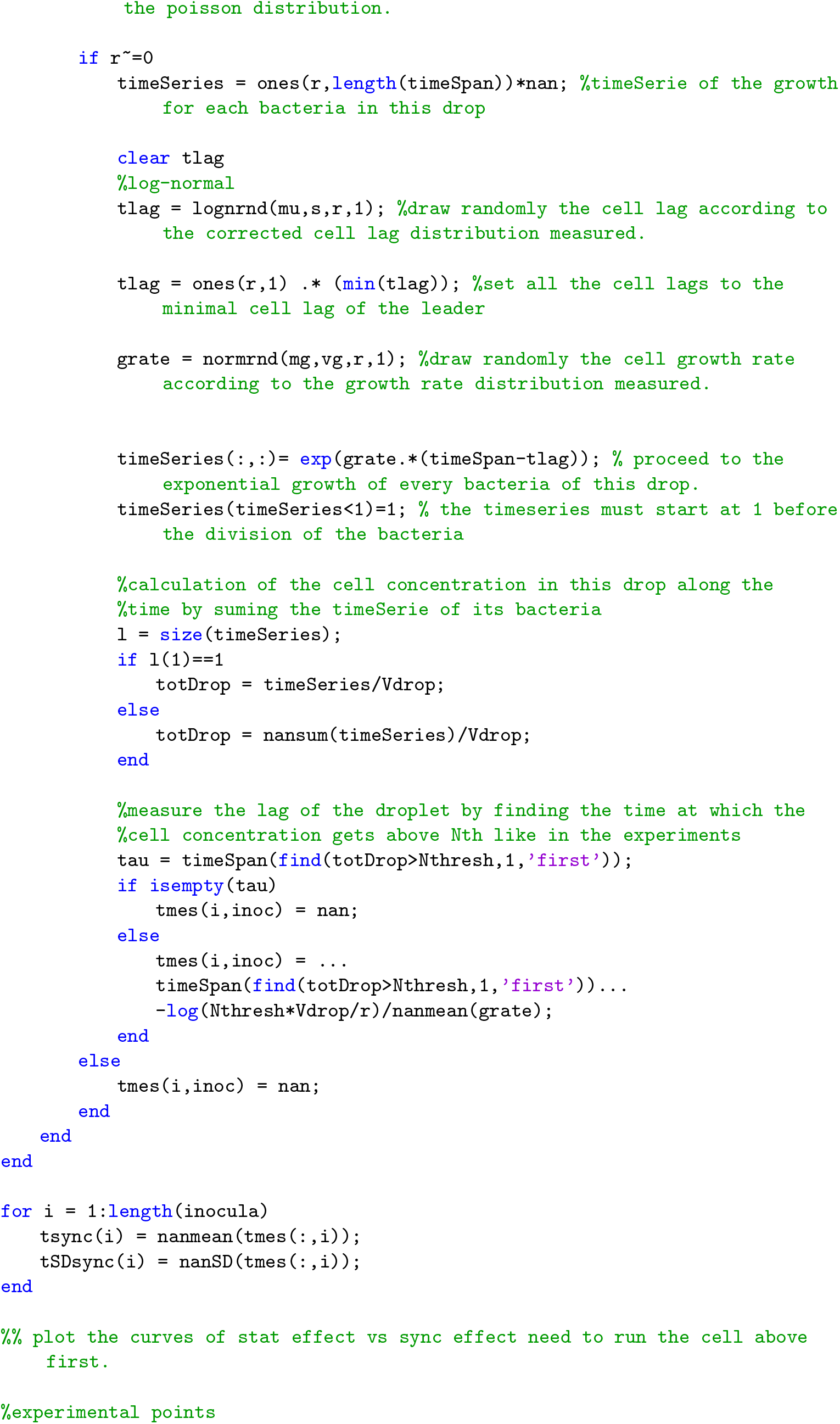
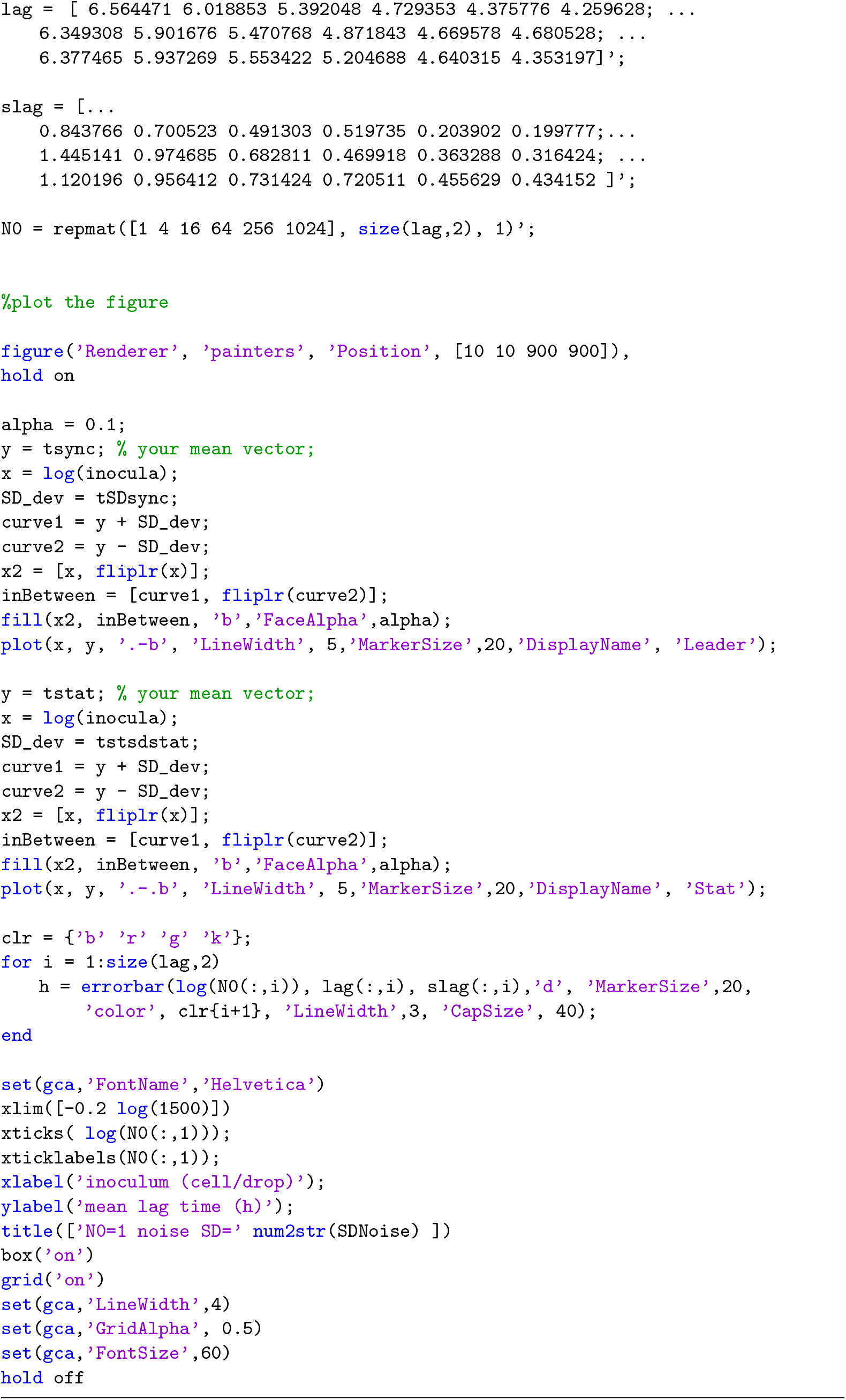
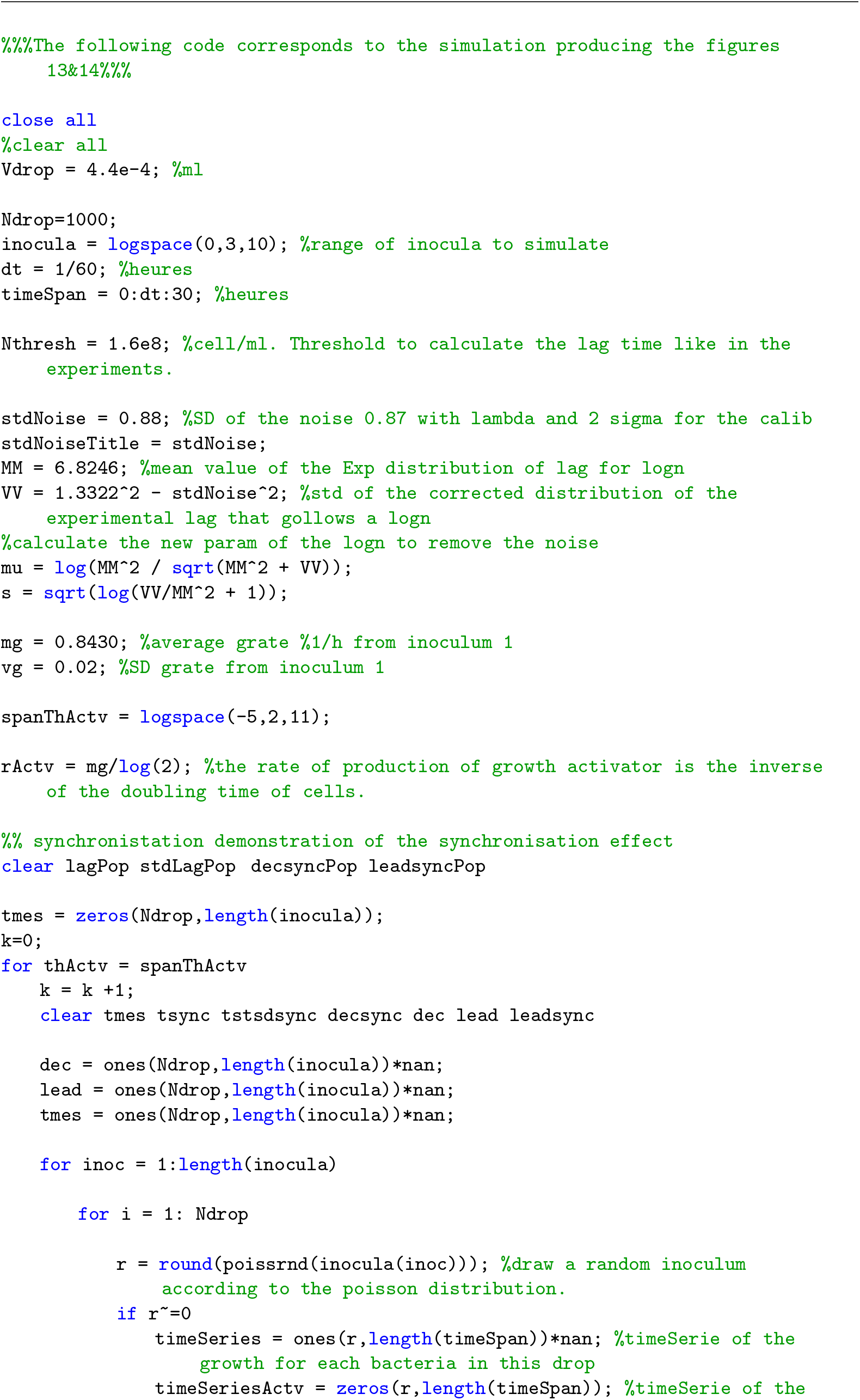
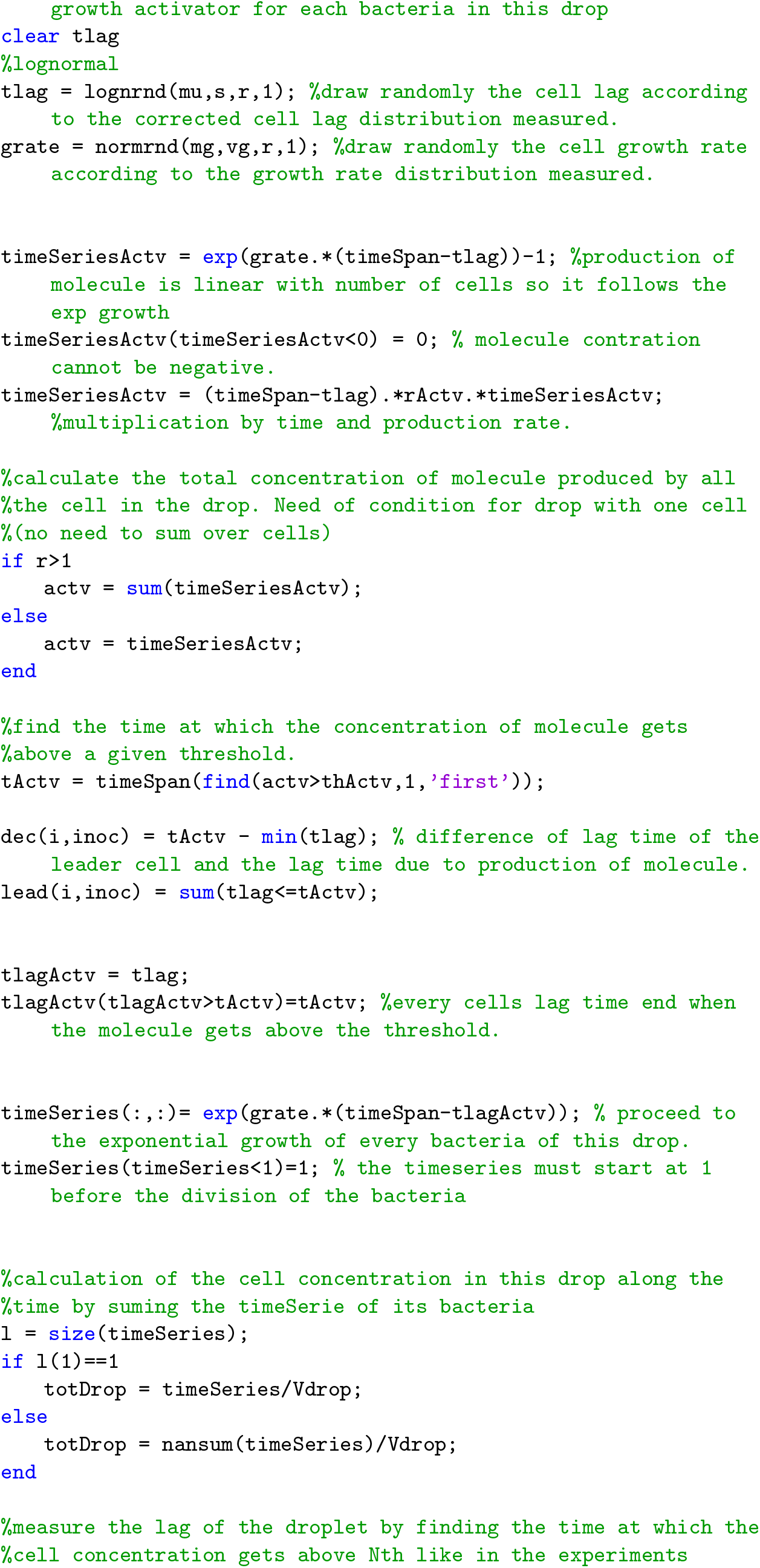
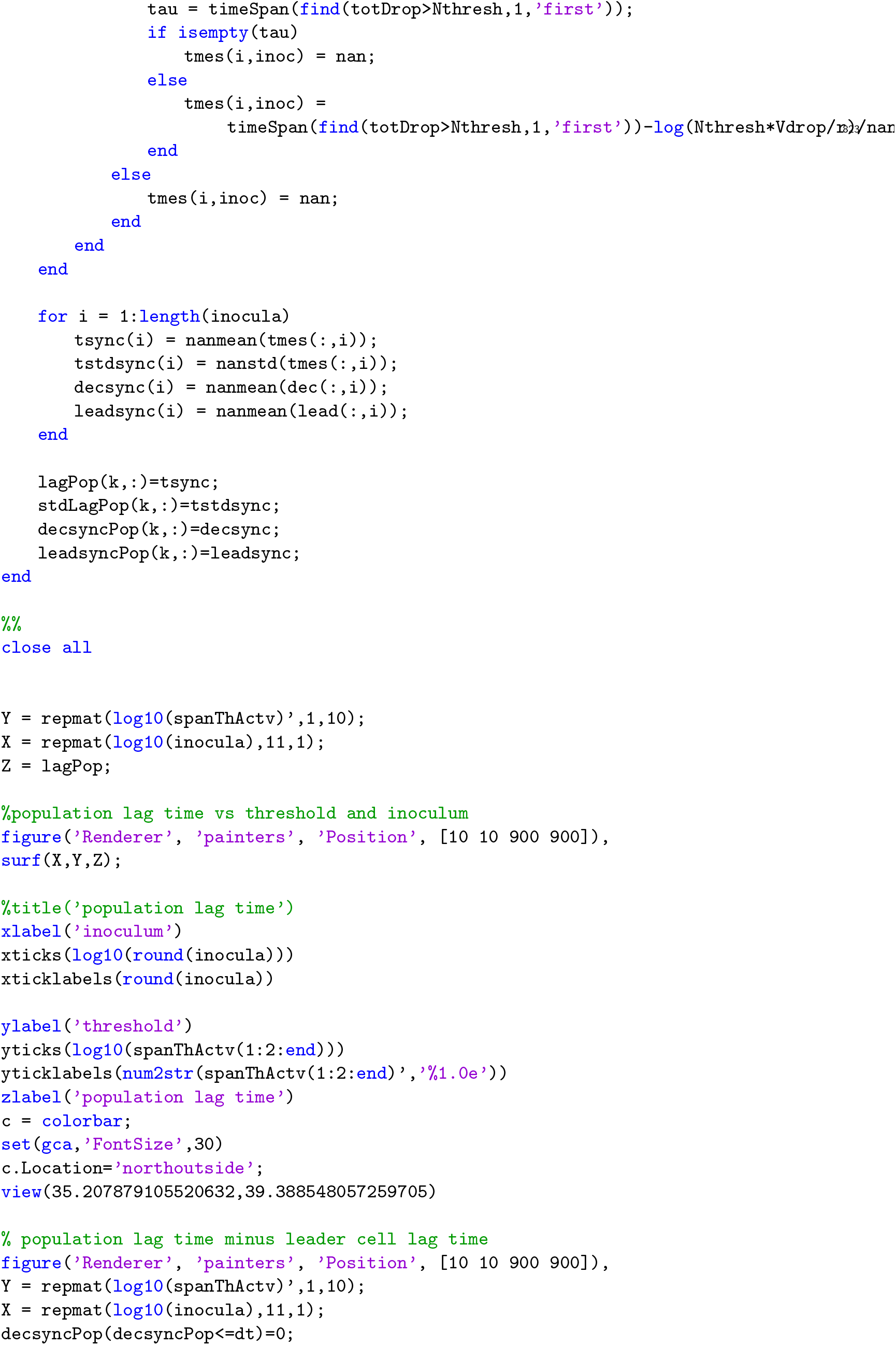
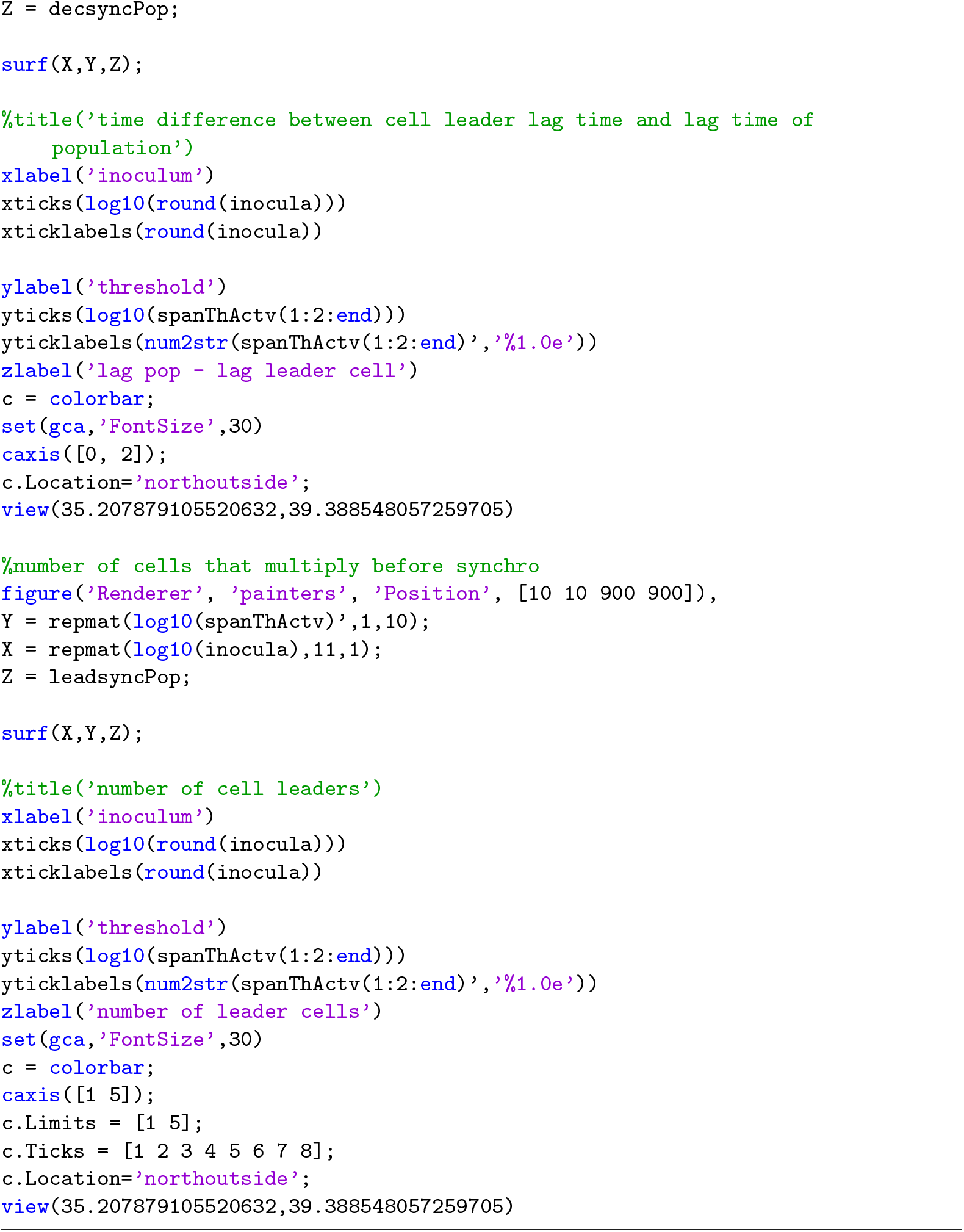

### Statistical properties for extreme values

Extreme Value Theory tells us that the minima of finite samples, drawn from some parent distribution, exhibit predictable statistical properties in the limit of large sample sizes [8]. These properties are testable experimentally in our system, where we can control the sample size - inoculum size *N*_0_, and repeat the sampling many times. The theory predicts how the mean and variance of sample minima depend on *N*_0_, given the parent distribution. It moreover contains the powerful statement that the distribution shape converges to a universal one in the limit *N*_0_ → ∞, which reflects the decay of the original distribution tails. Similar statements hold for maximum as well as other extreme values (e.g. second largest, etc).

To develop some intuition for the scaling of moments with sample size, imagine drawing *N*_0_ random variables from a normal Gaussian distribution. To estimate the minimal drawn number, we divide the real line into *N*_0_ equal-probability bins (see illustration below). On average, there will be one number drawn from each bin; therefore we may estimate that the minimal value lies within the lowest bin, in the range (−∞, *x*_1_). Clearly, the larger is the sample, the more bins we can use and still have an average of one number in the lower one; as the sample becomes large, we are increasing our chance of sampling low-probability events in the tail of the distribution.

To get the qualitative nature of the scaling, we seek a relation between the sample size *N*_0_, also the number of bins, and the values of the continuous variables *x* in the lowest bin. Comparing the probabilities,

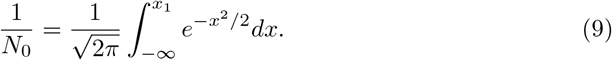

For large *N*_0_, *x*_1_ is very small and the integral is dominated by the uppermost limit. This means that if a random number is drawn from the lower bin, it is highly likely equal (or close to) the upper end of the bins; the chance of getting other values inside the bin are exponentially smaller. Therefore we approximate

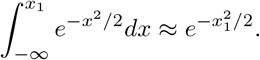

With this relation, we can solve for *x*_1_(*N*_0_) in Eq. 9 to find

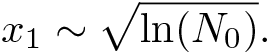

This value of *x*_1_ gives the scaling behavior of the minimal random number out of a large sample *N*_0_, on average. Computing the pre-factors correctly and estimating the sub-dominant terms requires more careful approximations.

Although this argument is made for Gaussian random variables, the same scaling holds also for the lognormal ones. Additionally, it was made for a zero-mean and unit-SD distribution; to account for arbitrary mean and variance of the parent distribution, in Fig. 3 we shift and scale by the nonzero mean and non-unit SD, respectively.

### Empirical testing of scaling property

A unique feature of our experiment is the measurement of population lag time distributions for different controlled inoculum size. Thus, we can test whether these distributions have a shape that is predicted by the EVT because the population lag times would be actually equal to a minimal cell lag times. In the following we show that all the distributions collapse on one another after appropriate scaling indicating that they have indeed a common shape underpinned by EVT. The analysis we propose can be done without knowing the parameters of the distribution and is a more robust approach compared to analysis made by using a fit which can be sensitive.

The most general form of the extreme value distribution was specified in Eq. 8 where *θ*_0_ is the location parameter, *γ* the scale parameter and *k* the shape parameter. The mean and variance of this distribution generally depend on these parameters. In particular, they are sensitive to *k*, for *k* = 0 the extreme value distribution is a Gumbel distribution and for *k* ≠ 0 it is a Frechet or Weibull distribution. Taking a practical perspective, we show below that it is not required to fit the parameters of the distribution in order to test for their scaling property; it is sufficient to empirically subtract the average and divide by the standard deviation.

For all cases where the first two moments exist, they are

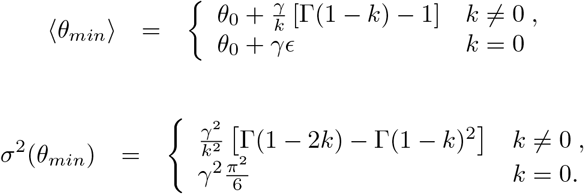

Where *ϵ* is the Euler–Mascheroni constant and Γ the gamma function. Although these are cumbersome expressions, they have the simple form

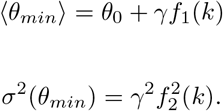

Therefore, the two-parameter scaling by the two first moments amounts to an affine transformation of the random variable:

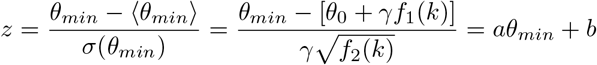

with *a, b* constants that can depend on *k*. It is not difficult to show (see section below) that if a variable is distributed according to a Generalized Extreme Value distribution (GEV) with shape parameter *k*, *θ_min_* ~ *GEV* (*θ*_0_, *γ, k*), then affine-transformed variables *z* = *a · θ_min_* + *b* are also GEV-distributed, with modified scale and shift parameters but with the same shape parameter: 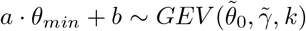. This means that the GEV distribution shape is invariant under affine transformations. Therefore, the distribution collapse of sampled data after empirical scaling by mean and SD provides a test for their common shape, and thus for their consistency with the extreme value properties.

Using the general expressions for mean and variance, we may express *σ*(*θ_min_*) as a function of ⟨*θ_min_*⟩

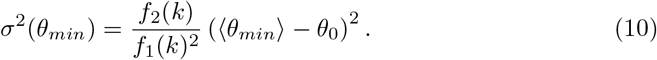

This relation was also tested in the measured data, as is depicted in the inset of the Fig. 3B.

### Invariance of GEV under affine transformation

We here show the invariance of distribution shape under affine transformation, for the entire GEV family of distributions. Suppose *x ~ GEV* (*x*_0_, *γ, k*) where *x*_0_ is the centering parameter, *γ* the scaling parameter and *k* the shape parameter. The cumulative form of the GEV is given by:

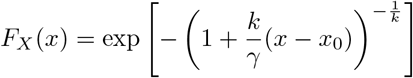

Now consider the variable *y* = *ax* + *b*: the cumulative form of the distribution of *y*(*F_Y_* (*y*)) can be derived from the distribution of *x* (*F_X_* (*x*)):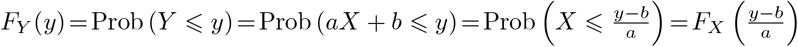. Therefore using the expression of *F_X_* we can find *F_Y_*. Using y_0_ = ax_0_+b,

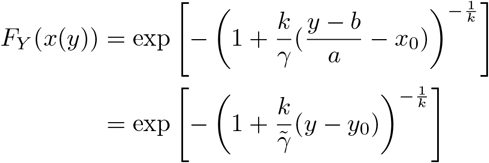

where 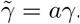. This shows that *y* is also distributed according to a GEV distribution, with modified shift and scale parameters but with the same shape parameter *k*.

It can be seen directly from the general expression of the mean and variance of the GEV family, that the scaled variable

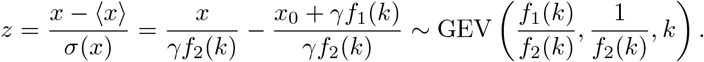

This GEV distribution of *z* has a mean ⟨*z*⟩ = 0, a standard deviation *σ*(*x*) = 1 and only a single parameter *k*. This property of extreme values distribution is used in the main text Fig. 3D to demonstrate that the population lag times follows the EVT and therefore that the population lag time is equal to the leader cell lag time.

**Figure 4.**
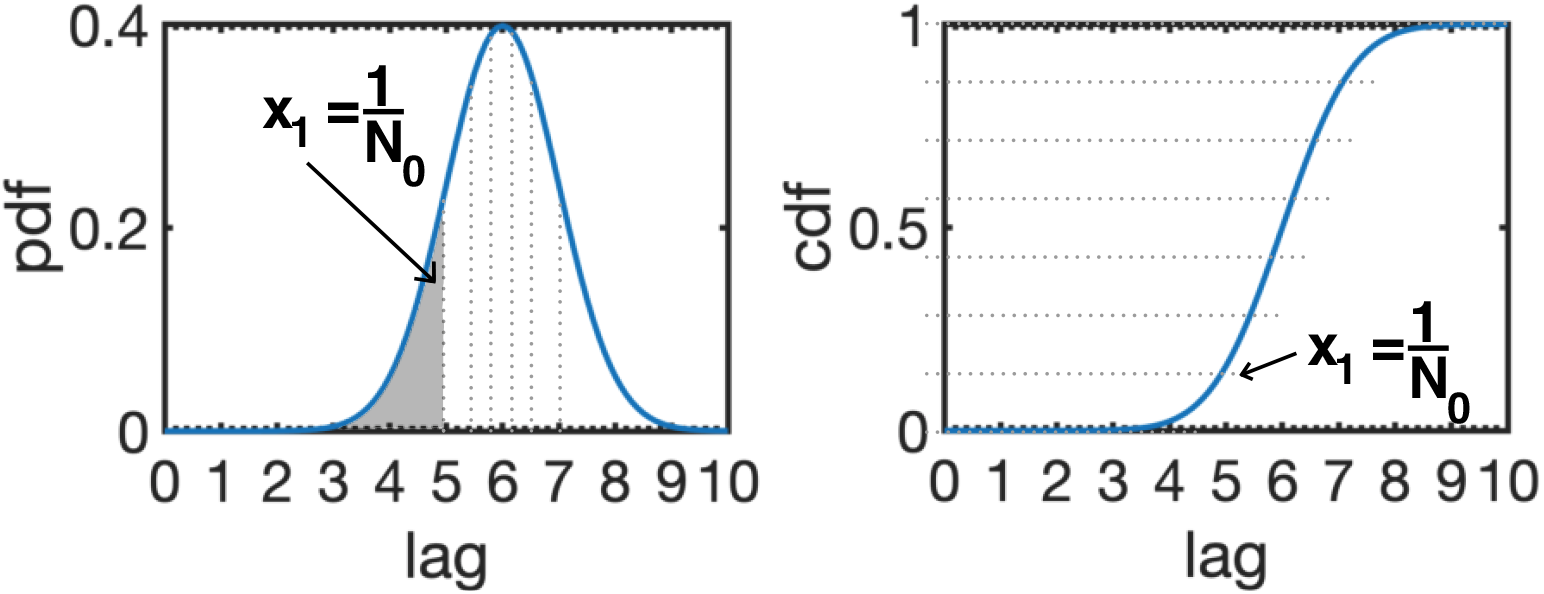
Drawing a finite sample of size *N*_0_ from a continuous Gaussian distribution. To estimate the minimal value drawn out of *N*_0_ values, we divide the real line into *N*_0_ equi-probable bins; in the figure, *N*_0_ = 7. The √estimate that the minimum resides in the lowest bin can be used to derive the scaling 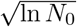 in the limit of large *N*_0_. Left we represent the probability density function of a Gaussian (mean 6h, SD 1h). The probability to have value in the lowest bin between 0 and 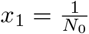 is equal to the area of this bin highlighted in grey. Right we represent the cumulative density function of the same Gaussian. The probability to have a value below or equal to 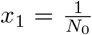 is equal to the y-value of the lower dotted line pointed out by the arrow.

**Figure 5.**
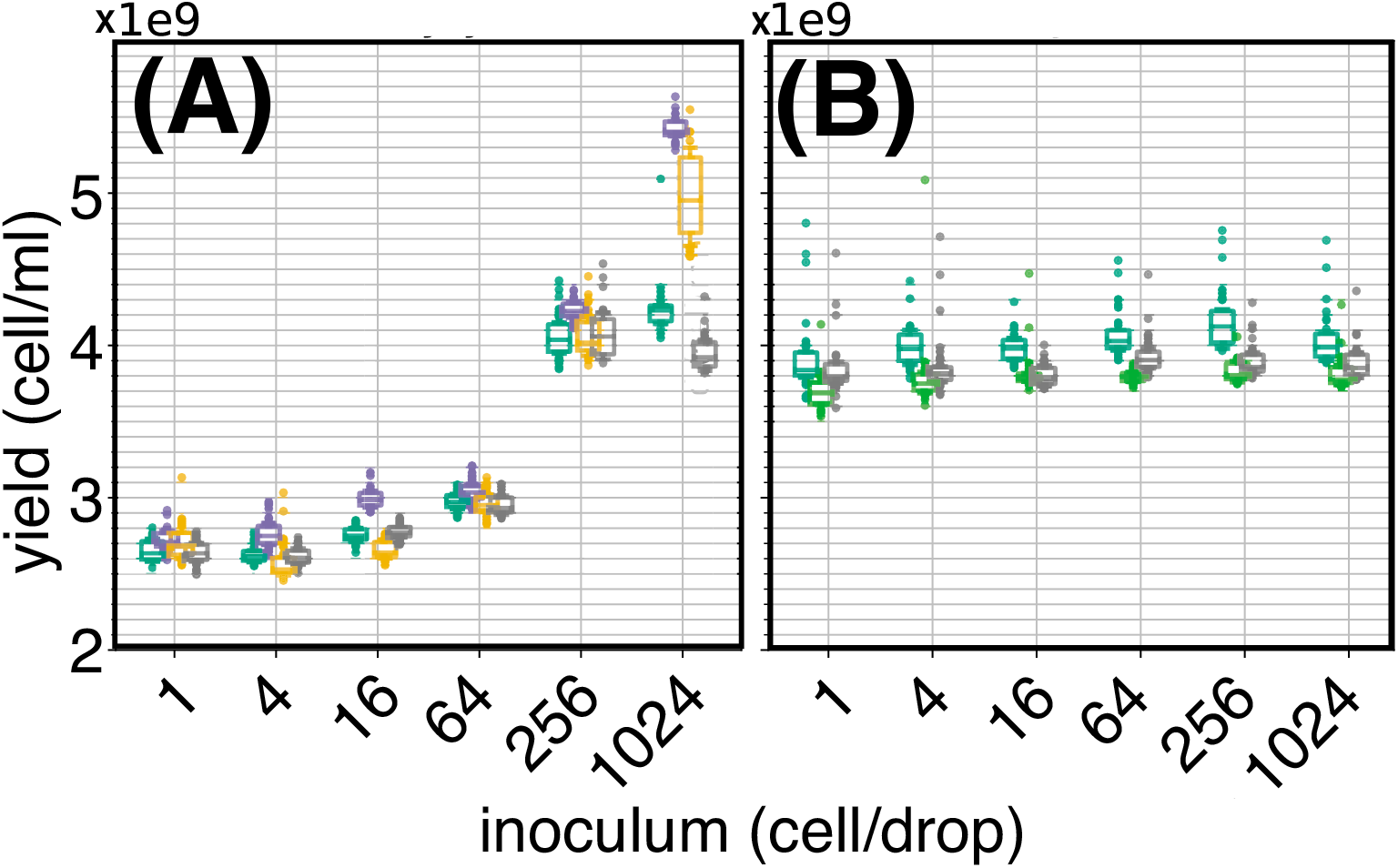
The maximal density of bacteria in the droplets depends on the abiotic inoculum. We measure the maximal density of bacteria reached for 6 different inoculum (1, 4, 16, 64, 256, 1,024 cells) in rich medium. Each box plot is calculated from 40 droplets (30 for the inoculum 1,024). The different colours correspond to independent replicates. **(A)** shows the yield of droplets prepared from a 100μl glycerol stock with a “naive” serial dilution that does not compensate the glycerol concentration across bacterial inocula. We observe that the maximal density of bacteria depends on the inoculum. At high dilution, *ie* small bacterial inocula, the maximal density reaches 2.6 · 10^9^ cell·ml^−1^ whereas at low dilution, *ie* large inoculum, the maximal density goes above 4 · 10^9^ cell·ml^−1^. (A) and (B) share the same y-axis. **(B)** shows the maximal density of an “aware” serial dilution such that the density of glycerol is kept constant across inocula. To do so the glycerol density is balanced by addition of an appropriate volume of glycerol (stock at 60% w/w) in the inocula. We see that balancing the glycerol in the droplets results in a constant maximal density of bacteria, whatever the inoculum of bacteria in the droplets. Thus, traces of glycerol coming from the frozen stocks influence the maximal density of bacteria. Diluting “naively” the glycerol of the frozen stock by 70x (together with the cells) yield an increase of 150% of the maximal bacterial density reached in droplets compare to a dilution of 70,000x. Thence, The millifluidic technology has the sensitivity to measure precisely such abiotic effect. In our work we always took care of balancing the glycerol concentration in the culture to keep it constant across inocula.

**Figure 6.**
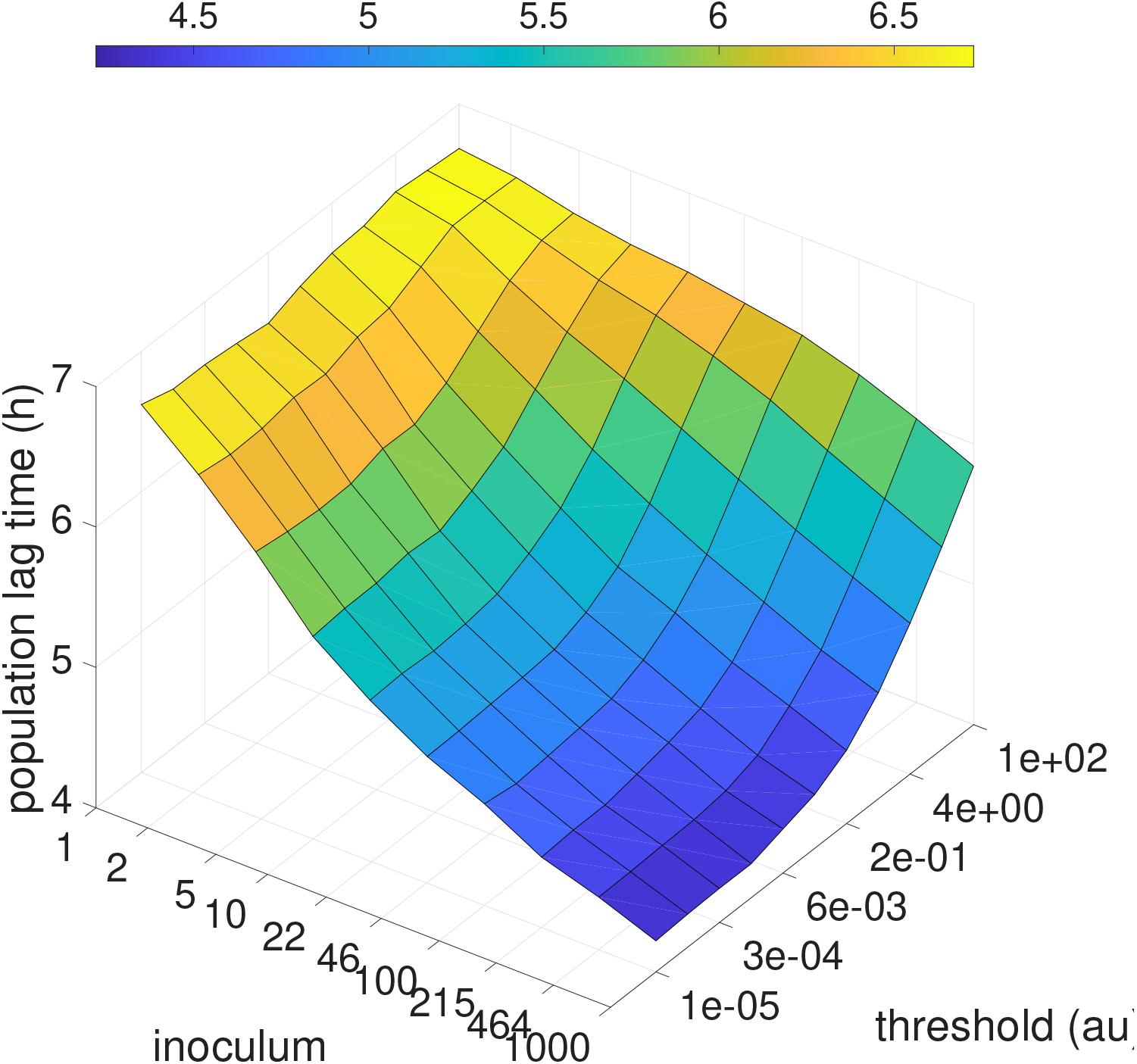
Population lag time in a model of growth activator production. Results of simulation where cells produce a growth activator, that accumulates to a threshold to trigger exit from lag phase and start of growth. The x-axis shows inoculum size in number of cells; the y-axis shows the threshold of the growth activator in arbitrary units. The z-axis and the color-map depict the resulting population lag time in hours. A significant inoculum effect - a dependence of population lag-time on inoculum size - is seen only at low activation threshold. Our experimental observations are in agreement with the smallest threshold value tested here, resulting in a strong dependence on the inoculum size.

**Figure 7.**
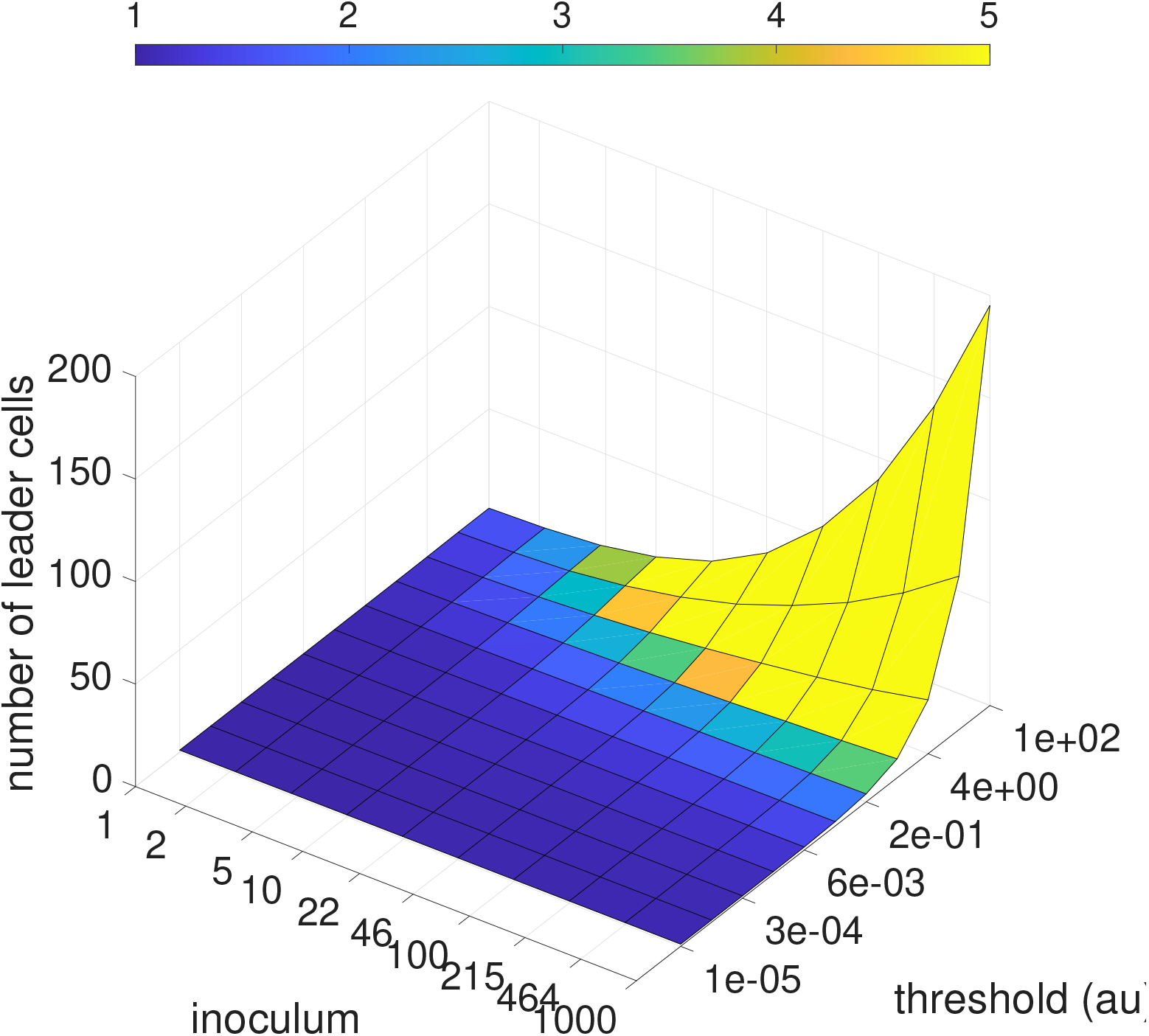
Number of leader cells in a model with growth activator production. For the same simulations than Fig. 6, shown is the number of cells which have exited their lag phase (according to their lag time value drawn from the single cell distribution) before the threshold was crossed. These cells can be considered as “leader cells”, because they have caused the others in the population to start growing in response to the cooperative signal, rather than according to their single cell lag time. The x-axis shows the inoculum size in number of cells. The y-axis shows the threshold of the growth activator in arbitrary unit. The z-axis and the color-map depict the number of leader cells. The color-map is saturated for a value of 5 cells (yellow) but eventually the number of leader cells go above 5 in a certain range of the plot. In the region relevant to our experiment, smallest threshold value, the number of leader cells is 1. In fact this is true in a large range of inoculum size and growth activator threshold. This is visible by the large region of the surface that remains dark blue.

**Figure 8.**
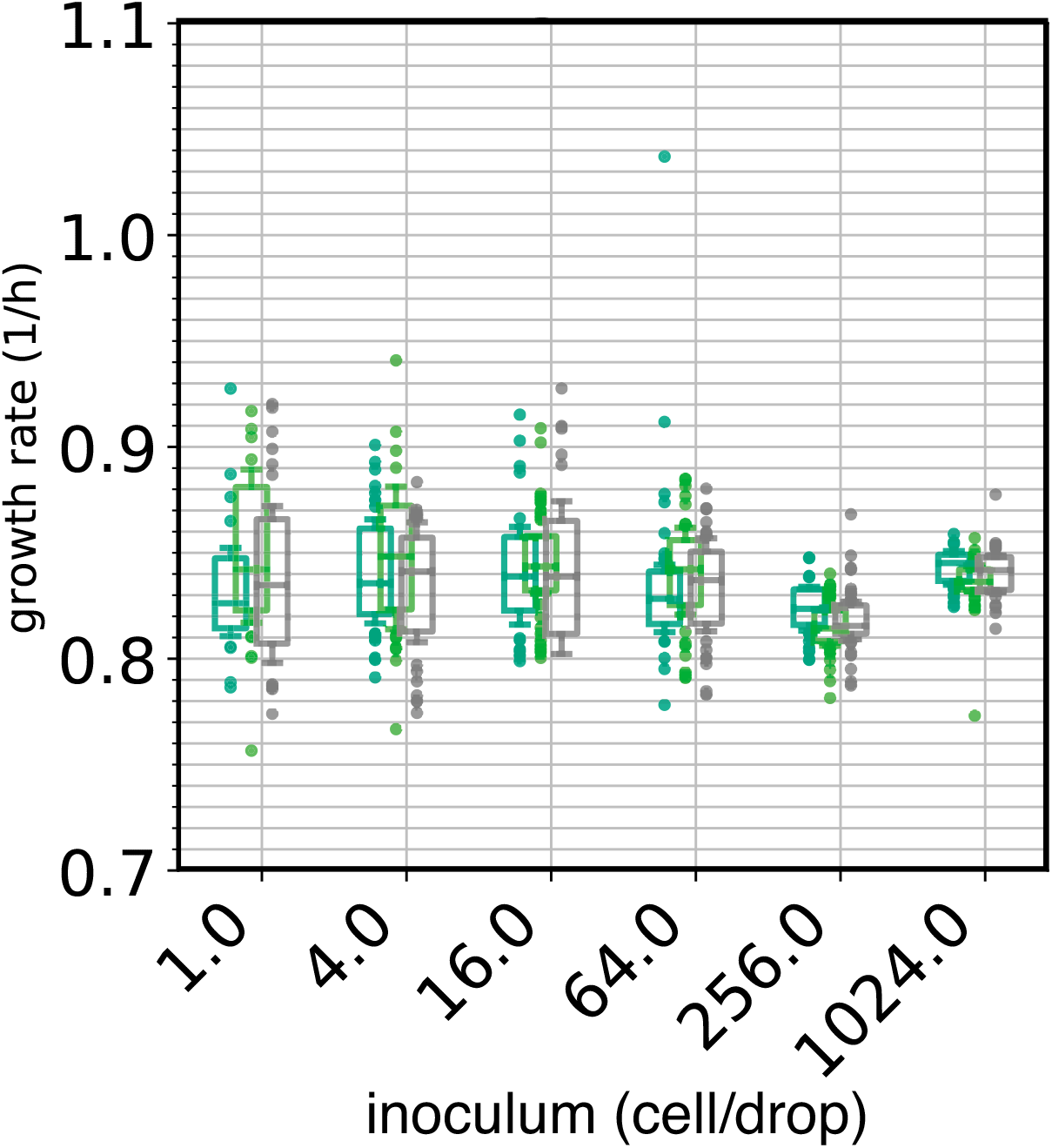
Growth rate in droplets does not depend on inoculum size. Boxplots of growth rate as a function of inoculum size estimated from time-series of 40 droplets (except 1,024 which is 30). The data are the same as in Fig 2A that shows the lag time of these droplets. The mean growth rate is approximately constant, with a median at 0.84 ± 0.02 *h^−^*^1^. The variance in growth rate decreases with inoculum size. The colors correspond to three independent experiments.

**Figure 9.**
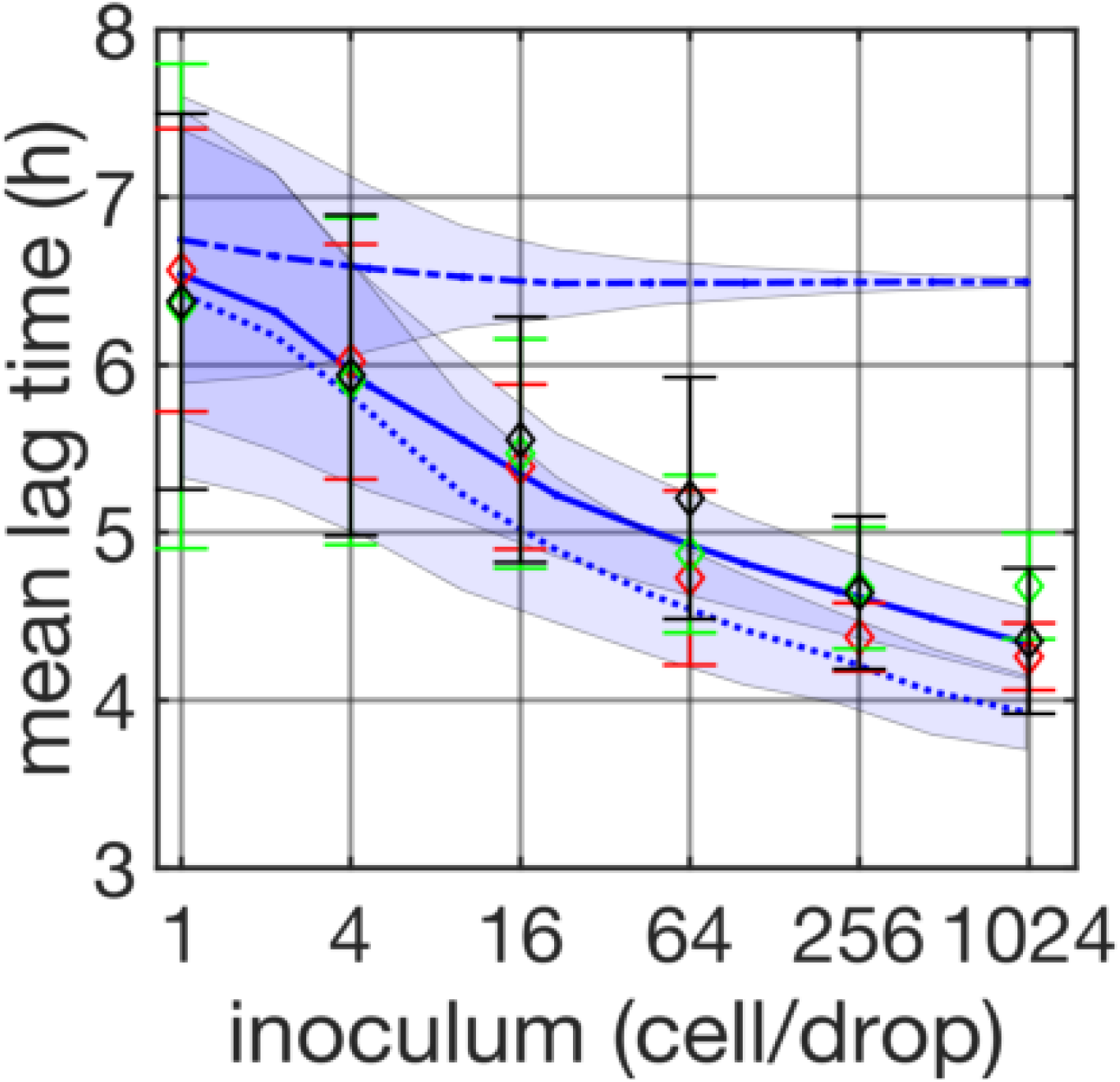
Effect of deconvolving the measurement error on single-cell lag time distribution. The same data shown in Fig. 2A are shown here. Results of the simulation where no deconvolution is done on the cell lag time distribution Fig. 3B is shown in dotted line. This simulation uses directly the raw cell lag time. It under-estimates significantly the mean lag time as a function of inoculum size.

**Figure 10.**
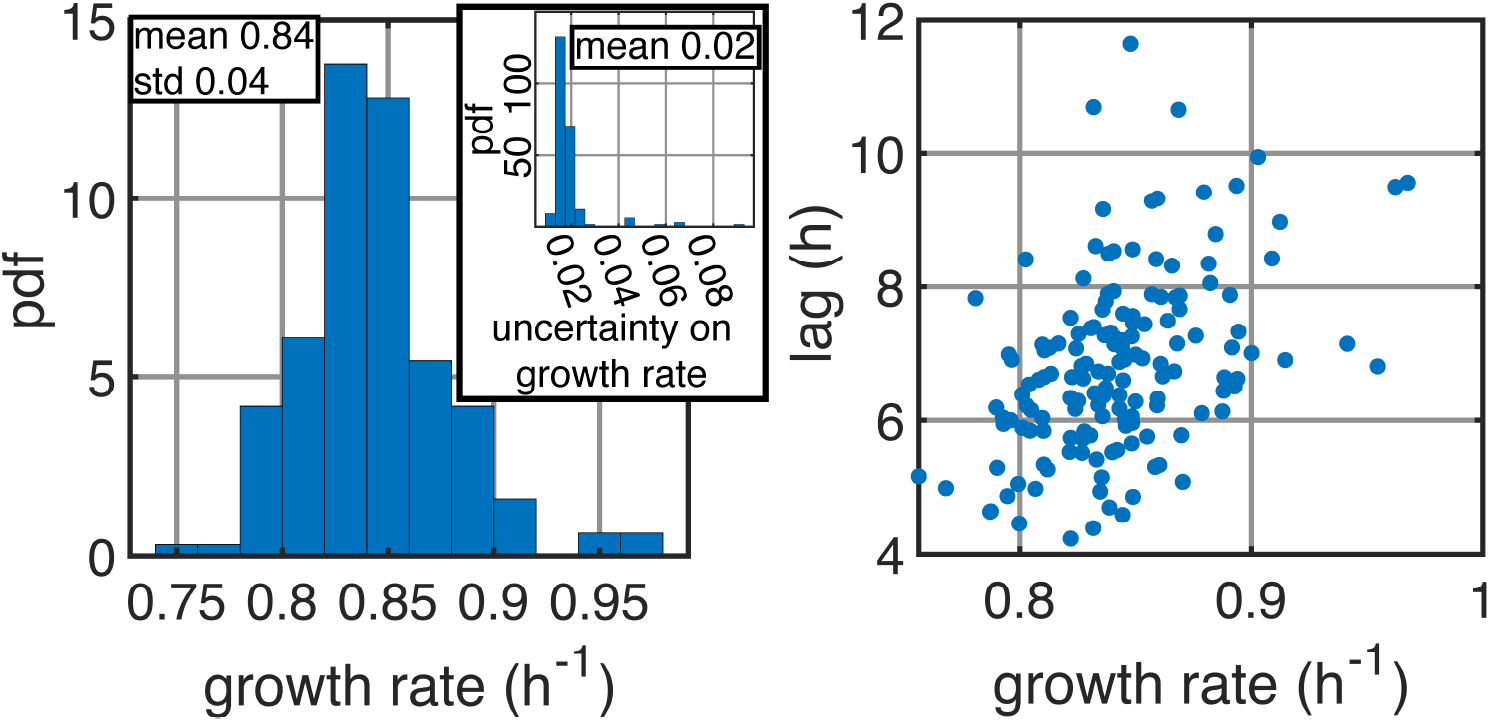
Statistical properties of experiment with inoculum 1. The left panel shows the probability density function of the growth rate in the experiment with inoculum 1 (Fig. 3B). The area of each bar is the number of observations in the bin. The growth rate is defined as the maximum value of the derivative of the growth curves (Fig. 1C). The estimation of the maximal value of the derivative for a droplet is given with its SD (shaded area around the derivative Fig. 1C). The inset reports the histogram of the SDs. The mean value of the SDs is taken as the typical error of the growth rate Δ*λ* = 0.02. The right panel shows that the lag time and the growth rate of each droplets is weakly correlated across (Pearson coefficient of 0.43). Every points correspond to one droplet of the experiment.

**Figure 11.**
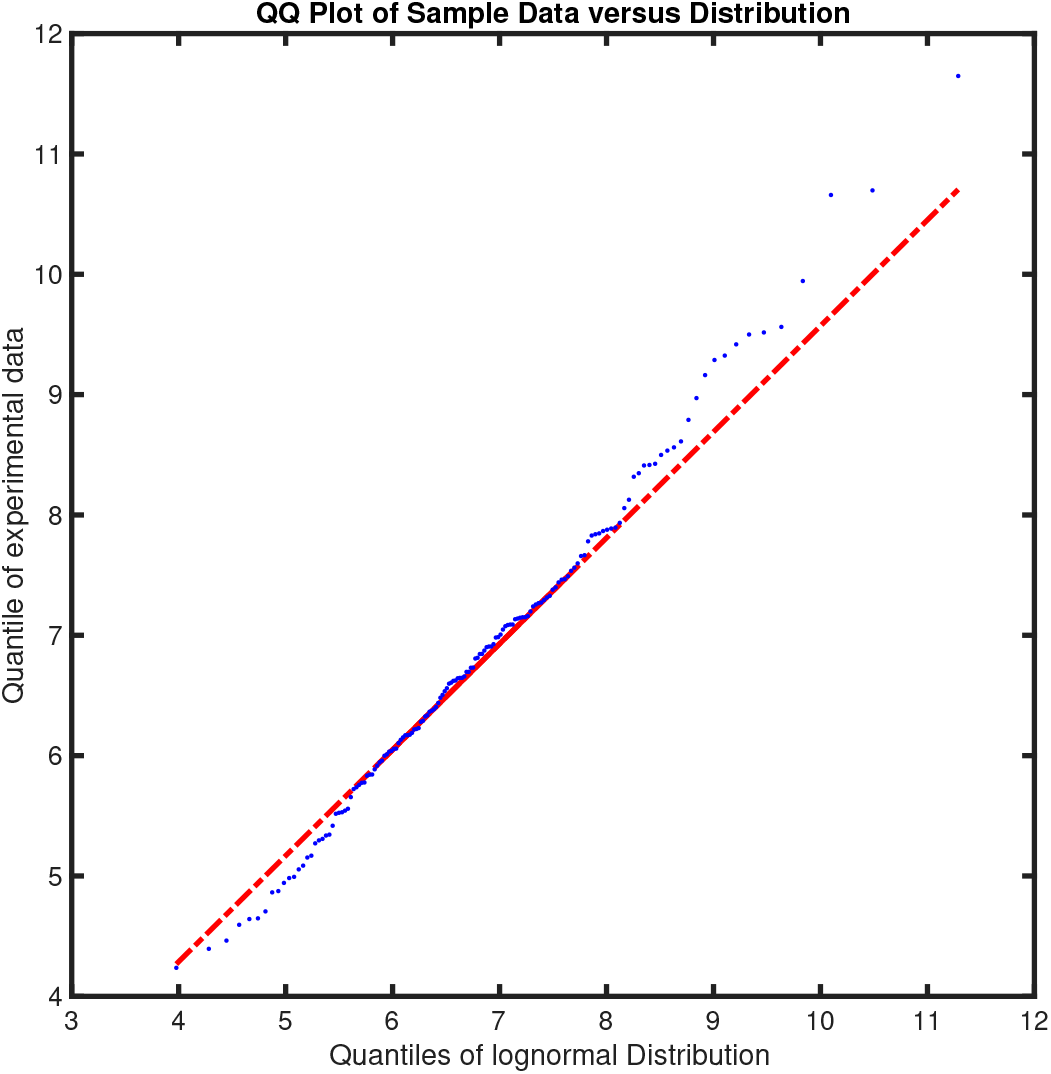
Quantile to quantile plot (qqplot) of cell lag time. To determine the distribution that underpin the cell lag time (*θ*) Fig. 2B we plot the quantile of a log-normam distribution versus the quantile of the distribution of the experimental measurements. The resulting quantile to quantile plot is well fitted by a line of slope 1 indicating that the experimental values follows a log-normal distribution.

**Figure 12.**
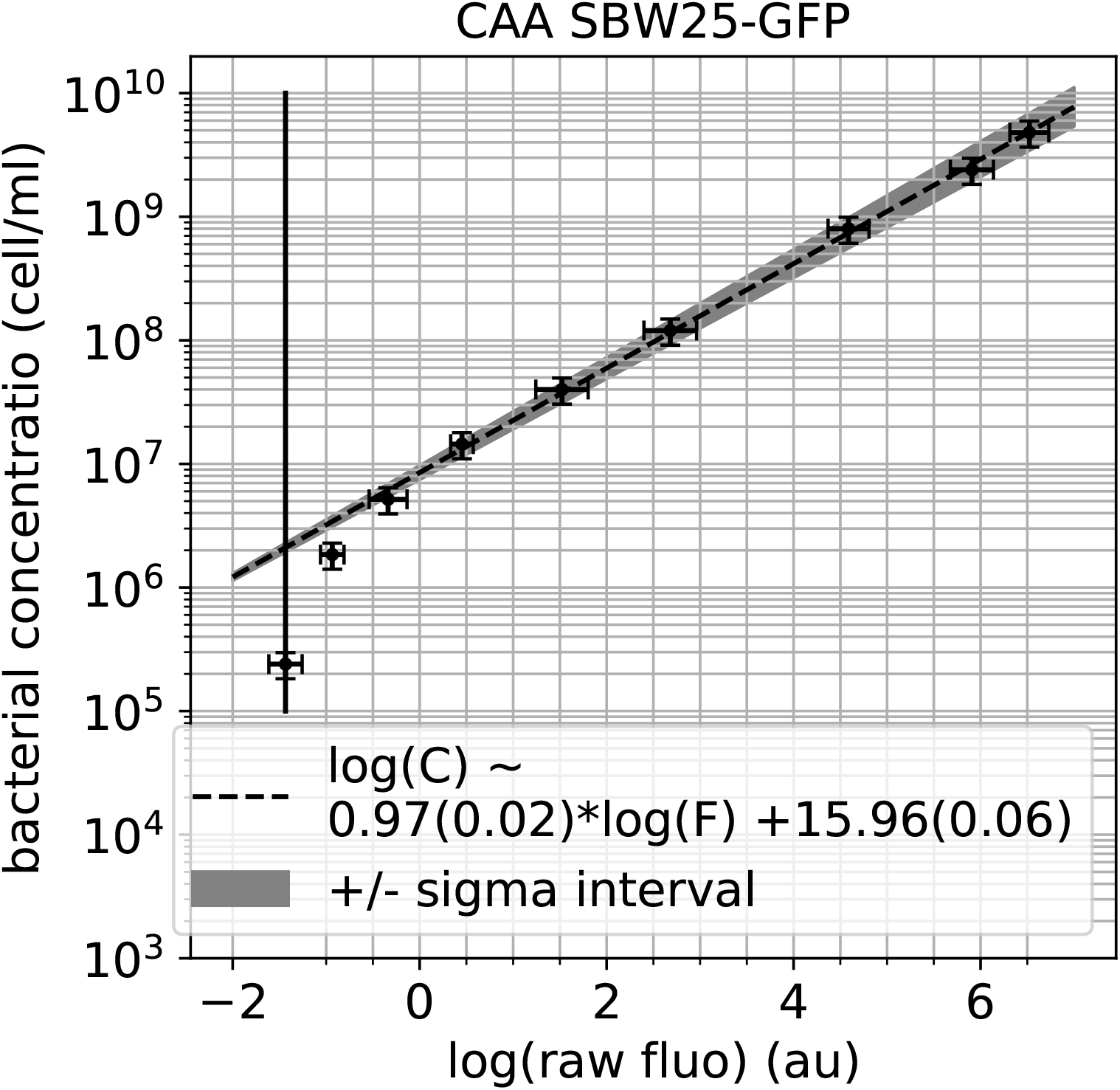
**Calibration curve** of the voltage given by the fluorescent detector in the channel GFP, called “raw fluo”, to a bacterial concentration in cell·ml^−1^. Every points correspond to 10 droplets in the millifluidic setup made with a given bacterial concentration. The range of bacterial concentration is achieved by diluting a mother culture grown over-night in CAA. The SD of the “raw fluo” across the 10 droplets is used as x-errobar. The bacterial concentration of the mother culture is measured with the standard protocol of serial dilution and count on agar plate. The agar-plate count is made with 10 replications allowing to estimate a SD of the mother culture counts. This SD is used in the plots as y-errorbar. The dashed line depicts a linear fit on the experimental points arranged on a log-log scale. Its equation is given in the inset with uncertainty. The vertical line depict the value of raw fluorescence for pure CAA medium (the blank).

**Figure 13.**
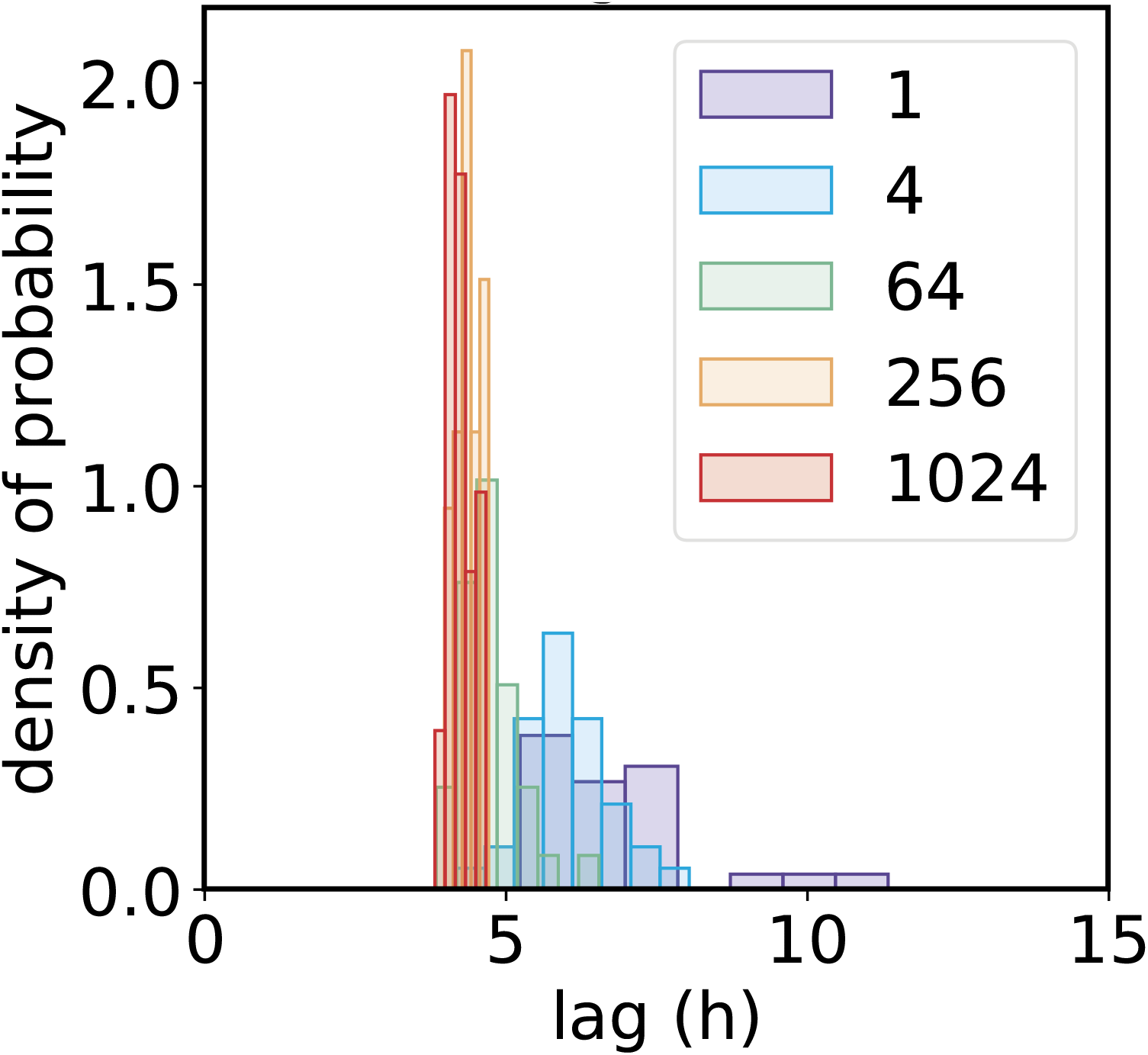
Histograms of lag time for a range of inoculum. It is the same data than one replica shown Fig. 3A but displaying the full histograms instead of points. The colors of the histograms displayed in legend indicate their corresponding inoculum (in cells/drop).

**Figure 14.**
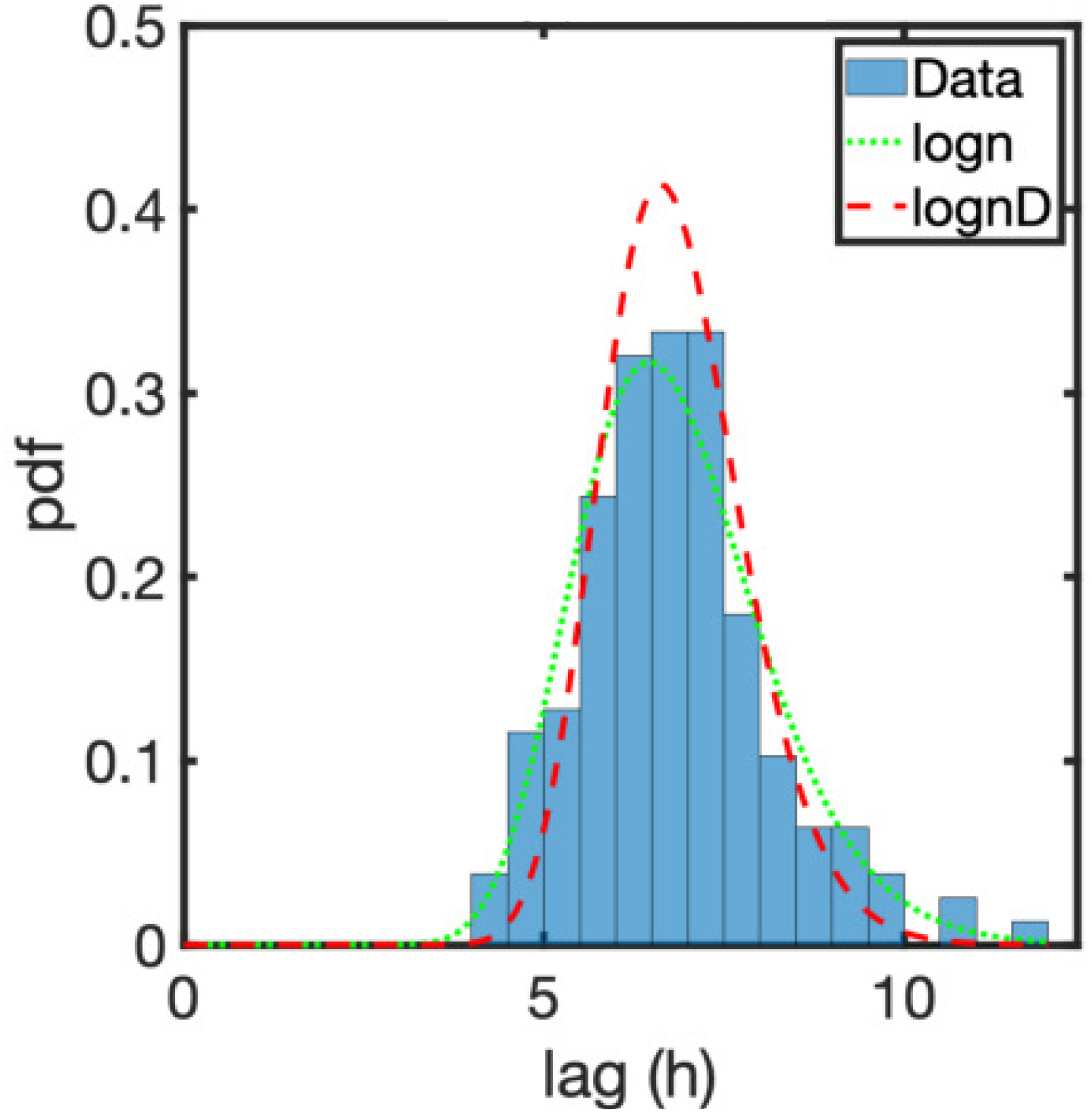
Probability density function estimates of the log-normal distribution of lag time. It the same data than Fig. 3B but in another way of representation often used. The “logn” in green dotted line is the fit on the data (blue bars). The “lognD” is the true log-normal distribution of lag after deconvolution of the gaussian noice.

**Figure 15.**
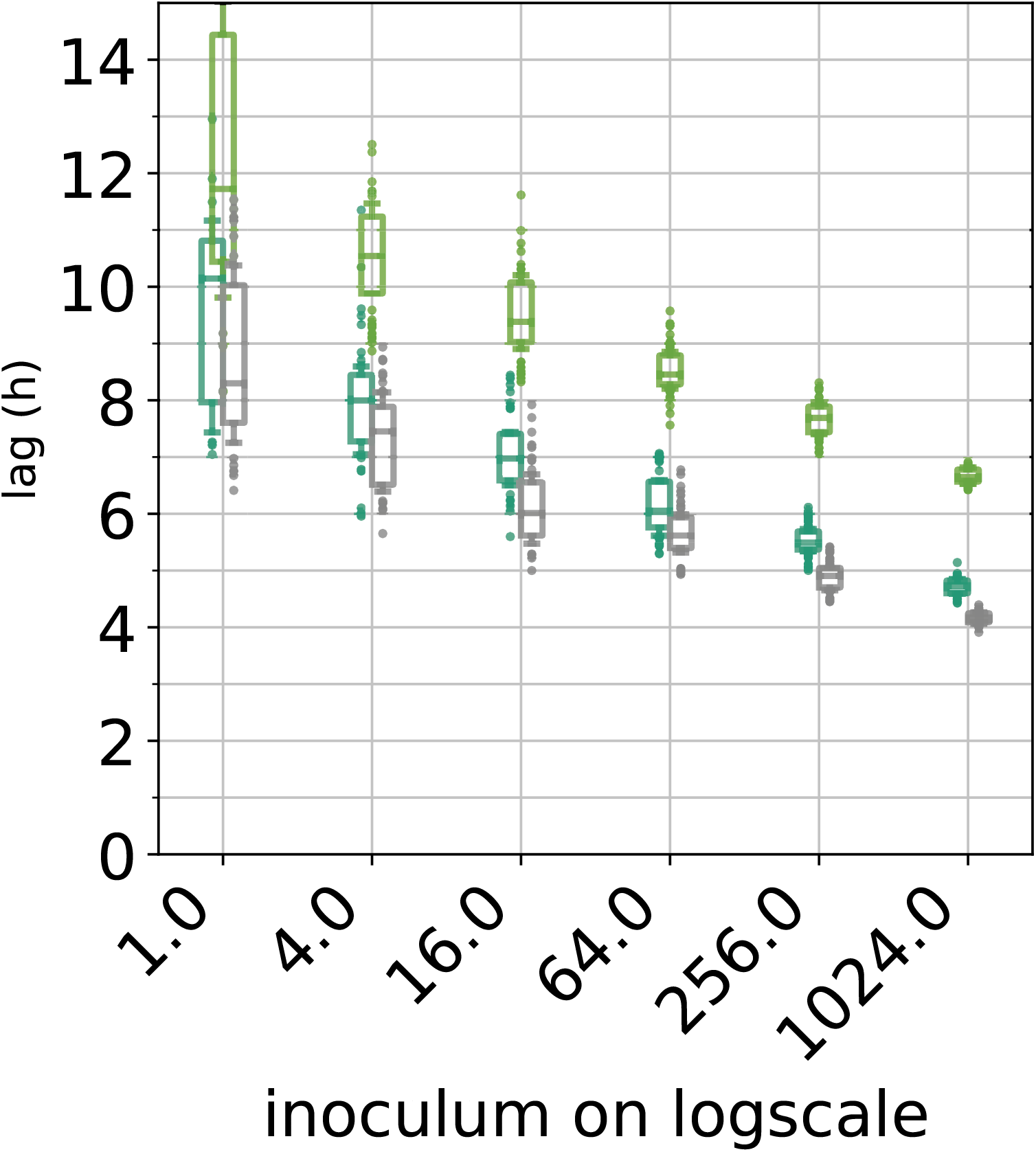
Inoculum effect on the lag for the strain *P. fluo* SBW25 PvdS229. This strain does not produce pyoverdine due to a mutation in *pvdS* a gene that encodes the extracytoplasmic family sigma factor *PvdS* and which directs expression of the pyoverdin biosynthetic genes (Cunliffe et al. 1995). The pyoverdin is an iron chelator the allows the pseudomonads to forage iron in their environment. The presence of the inoculum effect despite the non-production of pyoverdin indicate that this metabolite does not play a role in the coordinated exit of the lag phase. The three colors correspond to three independent experiments in fresh CAA. Data are represented in boxplot for every inoculum (with a little jitter for a better visualisation)

